# Quorum-sensing- and type VI secretion-mediated spatio-temporal cell death drives genetic diversity in *Vibrio cholerae*

**DOI:** 10.1101/2022.08.09.503374

**Authors:** Ameya A. Mashruwala, Boyang Qin, Bonnie L. Bassler

**Affiliations:** Department of Molecular Biology, Princeton University, Princeton, New Jersey 08544, USA.; Department of Mechanical and Aerospace Engineering, Princeton University, Princeton, New Jersey 08544, USA.; The Howard Hughes Medical Institute, Chevy Chase, MD 20815, USA.

## Abstract

Bacterial colonies composed of genetically identical individuals can diversify to yield variant cells with distinct genotypes. Variant outgrowth manifests as sectors. Here, we show that Type 6 secretion system (T6SS)-driven cell death in *Vibrio cholerae* colonies imposes a selective pressure for the emergence of variant strains that can evade T6SS-mediated killing. T6SS-mediated cell death occurs in two distinct spatio-temporal phases, and each phase is driven by a particular T6SS toxin. The first phase is regulated by quorum sensing and drives sectoring. The second phase does not require the T6SS-injection machinery. Variant *V. cholerae* strains isolated from colony sectors encode mutated quorum-sensing components that confer growth advantages by suppressing T6SS-killing activity while simultaneously boosting T6SS-killing defenses. Our findings show that the T6SS can eliminate sibling cells suggesting a role in intra-specific antagonism. We propose that quorum-sensing-controlled T6SS-driven killing promotes *V. cholerae* genetic diversity, including in natural habitats and during disease.

## Introduction

Bacteria track cell population density and the species composition of the vicinal community using a process called quorum sensing (QS). QS relies on the production, release, accumulation, and detection of extracellular signal molecules called autoinducers. QS enables groups of bacteria to synchronize gene expression, and collectively enact processes that demand many cells working together to make the task successful (Papenfort and Bassler, 2016; Waters and Bassler, 2005). In *Vibrio cholerae*, the causative agent of the cholera disease and the model bacterium used for the present work, two parallel QS pathways funnel information contained in autoinducers to a shared transcription factor called LuxO (Figure 1) (Miller et al., 2002). At low cell density (LCD), in the absence of autoinducers, the autoinducer receptors act as kinases ferrying phosphate to LuxO (Wei et al., 2012). LuxO∼P activates transcription of genes encoding four small RNAs (sRNA) called Qrr1-4. Qrr1-4 repress translation of HapR, encoding the master high cell density (HCD) QS regulator (Lenz et al., 2004). At HCD, when autoinducers have accumulated, the receptors act as phosphatases (Neiditch et al., 2005). LuxO is dephosphorylated and inactive. Production of the Qrr sRNAs is halted, HapR is translated, and it promotes expression of QS-controlled genes specifying collective behaviors (Lenz et al., 2004).

**Figure 1:**
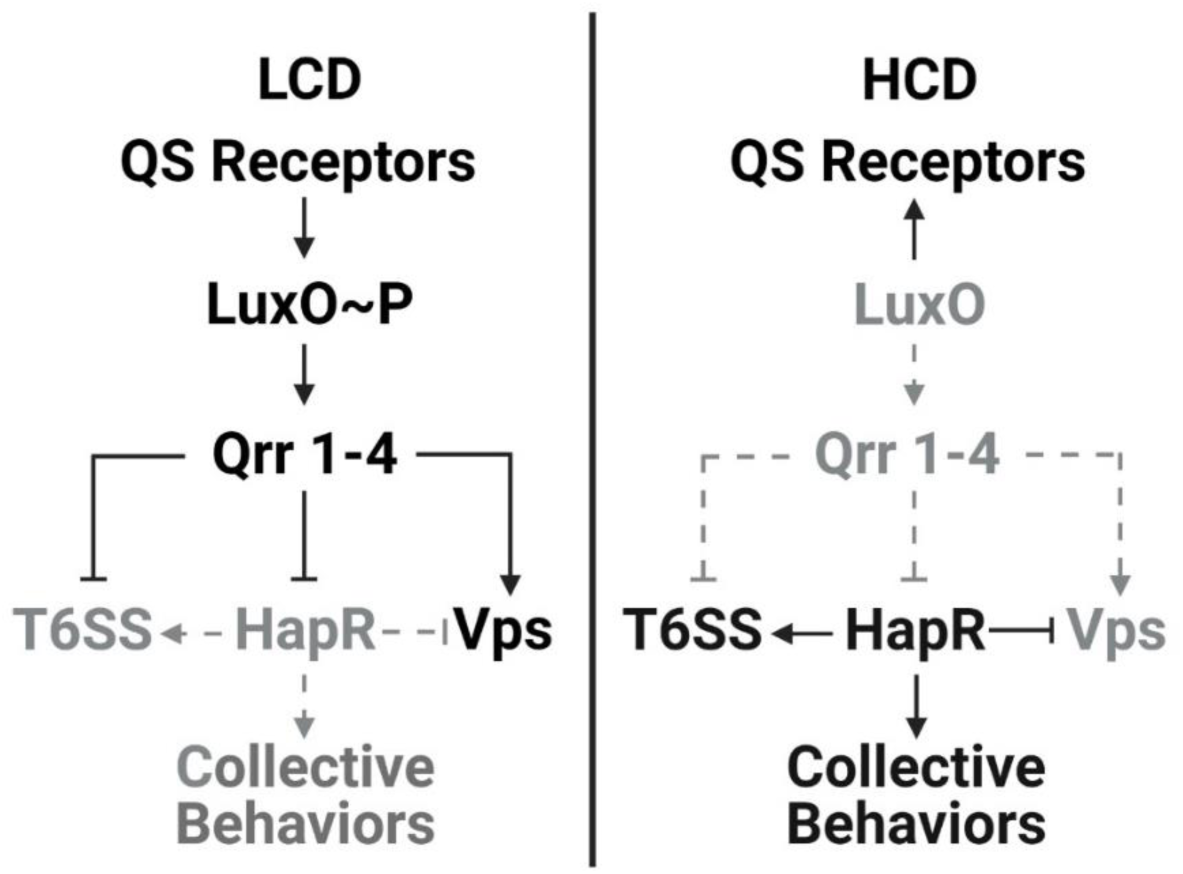
Simplified scheme for *V. cholerae* QS regulation of *t6ss* and *vps* genes. See text for details.

The *V. cholerae* type VI secretion system (T6SS) is a QS-regulated, contact-dependent protein delivery system that enables attack and elimination of competitor cells (MacIntyre et al., 2010; Pukatzki et al., 2006; Shao and Bassler, 2014). Briefly, T6SS structural components are assembled into a membrane-spanning spear-like device loaded with toxic effector proteins (Ho et al., 2014; Russell et al., 2014; Wang et al., 2019). The apparatus shoots the effectors into competitor cells by puncturing their cell walls. To prevent self-killing, T6SS-active bacteria produce immunity proteins that inactivate the toxic effector proteins (Ho et al., 2014; Russell et al., 2014; Wang et al., 2019). Other defenses, such as production of exopolysaccharide or capsular polysaccharide can also protect against incoming T6SS attacks (Flaugnatti et al., 2021; Hersch et al., 2020; Hood et al., 2010; Toska et al., 2018). In *V. cholerae*, *t6ss* genes are located in one large and three auxiliary clusters (Supplementary Figure 1) (Metzger et al., 2016). The large cluster encodes the proteins that make the type VI secretion complex and one effector-immunity protein pair. Each of the auxiliary clusters encodes one effector-immunity protein pair. The T6SS machinery is largely conserved among *V. cholerae* strains, however, its expression and regulation are strain specific. Important for this work is that a recently discovered cis-acting single nucleotide polymorphism (SNP) causes the El Tor environmental isolate 2740-80 to constitutively express its *t6ss* genes (Dörr et al., 2022; Ng et al., 2022). By contrast, the closely related pandemic isolate C6706 that lacks the cis-acting SNP does not express *t6ss* genes under laboratory conditions. In *V. cholerae* strains that do express *t6ss* genes, at LCD, the Qrr sRNAs repress *t6ss* gene expression by two mechanisms: they function directly to repress the large *t6ss* cluster and they function indirectly by repressing *hapR,* encoding HapR, an activator of the auxiliary *t6ss* gene clusters (Shao and Bassler, 2014). At HCD, when the Qrr sRNAs are not produced, the large *t6ss* gene cluster is derepressed and HapR activates expression of the auxiliary *t6ss* gene clusters. Direct cell to cell contact is required for T6SS toxin deployment (MacIntyre et al., 2010). Thus, restricting production of the T6SS machinery to high cell density could maximize killing efficiency.

Bacterial colonies are structured communities in which cells occupy micro-habitats with varying physical and chemical compositions (Bjedov et al., 2003; Hashuel and Ben-Yehuda, 2019; Saint-Ruf et al., 2014). These heterogenous environments impose distinct selective pressures for advantageous variant genotypes to arise. Often, the appearance of variants manifests as colony sectors. In *Staphylococcus aureus* and *V. cholerae*, such variants can display increased virulence in animal models and/or resistance to antibiotics (Finkelstein et al., 1992; Holmes et al., 1975; Koch et al., 2014; Servin-Massieu, 1961). Understanding the mechanisms driving colony variation could provide insight into the general emergence of new genotypes and their corresponding traits.

Here, we investigate the molecular mechanisms underlying colony sectoring in *V. cholerae.* We find that sectoring is preceded by T6SS-mediated cell death which occurs in two different spatio-temporal phases, each driven by distinct T6SS effectors. Tracking cell death using fluorescence microscopy shows that during the first phase, which occurs along the colony rim, T6SS-driven killing imposes a selective pressure for variant strains containing QS-inactivating mutations to arise. Loss of QS activity confers protection from T6SS-killing by two mechanisms. First, production of vibrio polysaccharide (VPS), normally a QS-repressed trait, increases and VPS acts to shield cells from incoming T6SS attacks. Second, elimination of QS-dependent activation of *t6ss* gene expression reduces overall T6SS-killing events. These changes confer regional growth advantages to the QS loss-of-function mutants which manifest as outgrowth into colony sectors. The second cell death phase, which occurs in the colony interior, while requiring a T6SS toxin, does not affect sectoring, and intriguingly, does not rely on the T6SS-injection apparatus. We propose that T6SS-driven intra-specific antagonism promotes *V. cholerae* genetic diversity, including in natural habitats and during disease, both of which are well known to select for *V. cholerae* variants that display reduced QS activity.

## Results

### *Vibrio cholerae* undergoes QS-dependent sectoring

Certain *Vibrionaceae* bacteria, including strains of *Vibrio vulnificus*, *Vibrio parahaemolyticus*, *Vibrio harveyi*, and *V. cholerae* form colonies that, over time, develop sectors that differ in opacity compared to the original colony (Chatzidaki-Livanis et al., 2006; Finkelstein et al., 1992; McCarter, 1998; Simon and Silverman, 1983). Presumably, other characteristics, not visible to the eye, are also regionally altered in the sectors and/or elsewhere in such colonies. The mechanism driving particular vibrios to form heterogeneous communities is not known. Here, we investigate the molecular underpinnings of colony sectoring using *V. cholerae* as our model system.

Following ∼2 days of incubation on solid LB medium, colonies of the O37 serogroup strain V52 and *V. cholerae* El Tor biotype strain C6706 did not sector and their morphologies remained uniformly translucent (Figure 2A). By contrast, the closely related *V. cholerae* El Tor biotype strain 2740-80 formed opaque sectors that were distinct from the translucent morphology of the initially growing colony (Figure 2A). We purified isolates from ten individual *V. cholerae* 2740-80 sectors. Each isolate formed a homogeneous opaque colony that did not sector, suggesting that these variants have acquired mutations that lock them into the phenotype of the sector. The levels of opacity differed between isolates, with the most extreme variant colonies displaying wrinkled morphologies (Figure 2A shows seven of these isolates). In *V. cholerae*, over-production of Vibrio polysaccharide (Vps) confers an opaque and wrinkled colony appearance, indicating that the variants from the sectors may have acquired mutations that drive increased Vps production. To investigate this possibility, we used whole genome and Sanger sequencing to successfully pinpoint the mutations that had occurred in nine of the isolated *V. cholerae* 2740-80 variants. Seven variants possessed a single nucleotide change, a deletion, an insertion, or an insertion-element aided interruption in genes encoding the *V. cholerae* master QS regulators LuxO (3 variants) and HapR (4 variants) (Supplementary Table 1 and designated in Figure 2A). One variant acquired a mutation in the 3’ UTR of the gene encoding the cold shock protein CspA, and the final variant had a mutation in *pyrG* encoding CTP synthase (Supplementary Table 1). Here, we focus on how alterations in QS drive changes in *V. cholerae* colony morphology and sectoring capability. We remark on the *cspA* and *pyrG* mutants in the Discussion.

**Figure 2:**
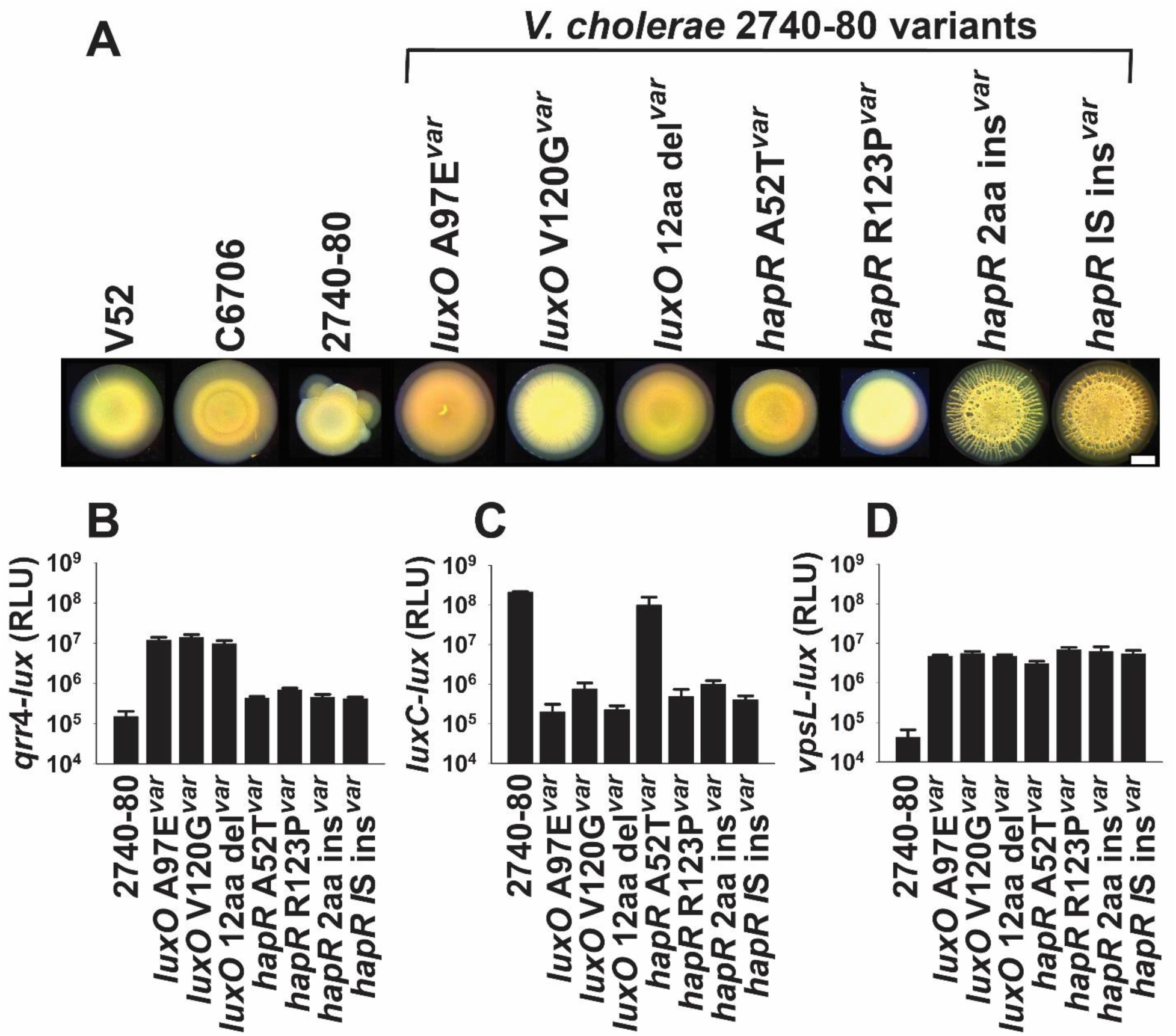
*V. cholerae* 2740-80 undergoes QS-dependent sectoring. (A). Brightfield stereo-microscope images of 2-day old colonies of the indicated *V. cholerae* strains. Scale bar, bottom right = 1 mM. (B-D). Transcriptional activities of (B) *qrr*4-*lux*, (C) *luxC-lux*, and (D) *vpsL*-*lux* in the indicated strains. Variant strains are annotated with the “var” suffix. In B-D, data represent average values from biological replicates (n=4), and error bars show SDs. Relative light units (RLU) are bioluminescence output divided by culture optical density.

To explore the connection between QS, sectoring, and colony morphology, we assessed whether the mutations in *luxO* and *hapR* that arose in the sectors specified gain- or loss-of-function alleles. Both LuxO and HapR are transcription factors. To measure their activities, we engineered luciferase (*lux*) transcriptional reporter fusions to well-characterized LuxO-controlled (*qrr*4) and HapR-controlled (*luxC*) promoters (designated *qrr*4-*lux* and *luxC-lux*, respectively). First, regarding LuxO: LuxO is phosphorylated and activates *qrr*4 transcription at LCD (Figure 1). The *V. cholerae* 2740-80 variants harboring mutations in *luxO* expressed ∼100-fold more *qrr*4*-lux* than did *V. cholerae* 2740-80 at HCD. This result shows that the *luxO* variants are gain-of-function alleles (Figure 2B). Regarding HapR: HapR is produced and functions at HCD, when it activates transcription of genes (Figure 1). All of the *hapR* variants except one expressed ∼100-1,000-fold less *luxC-lux* than did *V. cholerae* 2740-80 at HCD, showing that the HapR variants are either attenuated- or loss-of-function alleles (Figure 2C). Only the HapR A52T variant did not display an altered *luxC-lux* level (Figure 2C). HapR A52T has been studied previously. The A52T alteration affects HapR binding to DNA to different extents at different target promoters, but it does not affect binding to the *luxC* promoter (van Kessel et al., 2013). Thus, all but one of the QS mutants identified from the colony sectors “lock” the cells into the LCD QS mode.

QS promotes Vps production at LCD in *V. cholerae* (Hammer and Bassler, 2003). To connect the QS locked-LCD variant phenotypes to their opaque/wrinkled morphologies, we introduced a *vpsL-lux* transcriptional fusion into the strains and measured the output. *vpsL* encodes a biosynthetic protein required for Vps production. All of the LCD-locked QS variants displayed increased *vpsL-lux* expression compared to *V. cholerae* 2740-80 (Figure 2D). Thus, the LCD-locked QS states of the variants increases Vps production, and excess Vps converts the colonies from translucent to opaque/wrinkled. Importantly, while the colony wrinkling morphologies of the LCD-locked QS variants differed one from the other, none of the variant colonies sectored (Figure 2A). Thus, we infer that the HCD QS mode drives colony sectoring. Moreover, the sectors contain cells with genotypes that differ from that of the parent strain and the different mutations in the cells in the sectors underlie their distinct morphologies. Because the QS receptors funnel all sensory information to LuxO, and LuxO functions upstream of HapR in the cascade (Figure 1), in the remainder of this work, we focus on the *luxO* variant strains to understand how the LCD-locked state influences colony sectoring.

### T6SS-activity drives spatio-temporal cell death that precedes colony sectoring

*V. cholerae* 2740-80, which sectors, constitutively expresses the genes encoding its T6SS, while *V. cholerae* C6706, which is also an El Tor biotype strain but does not sector, does not express *t6ss* genes under laboratory conditions. *V. cholerae* uses Vps as a physical barrier to block T6SS attacks (Toska et al., 2018). In *V. cholerae* 2740-80, at LCD, LuxO∼P activates *vps* gene expression and represses *t6ss* gene expression (Figure 1) (Hammer and Bassler, 2003; Shao and Bassler, 2014). Thus, in our *V. cholerae* 2740-80 LCD-locked LuxO QS variants, *vps* expression is higher (Figure 2D). Similarly, at HCD, HapR represses *vps* expression, so in our *hapR* loss-of-function mutants, *vp*s expression increases (Figure 2D). Based on these patterns, we wondered whether cells in colonies of *V. cholerae* 2740-80 undergo T6SS-dependent killing. If so, individual cells that acquire mutations, such as in QS components that confer an increased ability to produce Vps, would reap growth advantages because they could use Vps to evade T6SS killing. This growth advantage could manifest in outgrowth as a sector. We designed experiments to test these ideas.

First, we assessed whether colonies of *V. cholerae* 2740-80 undergo T6SS-dependent killing and, if so, whether this affects colony sectoring. To do this, we quantified cell death in colonies of *V. cholerae* 2740-80 and in an isogenic strain lacking all four pairs of T6SS effector-immunity proteins (hereafter: Δ8 strain). We used time-lapse fluorescence microscopy to track live and dead cells and colony sectoring. For this analysis, all cells constitutively produced the mKO fluorescent protein (Red) to enable imaging of live cells and we used the fluorescent dye SytoX (Cyan) to mark dead cells. To enable visualization and fluorescence quantitation across the colonies, including in regions with sectors, we collapsed the time-series data into single images by generating projections across time. Representative time-series images are displayed in Figure 3A and Figure 3B,C shows the time-projections. To quantify spatio-temporal cell death in non-sectored regions, we reduced the time-lapse data into space-time kymographs (Figure 4). In both the time-projections and the kymographs, data were mapped using colors as quantitative readouts for intensities. There are many features in the images that differ between the strains under study. We focus on only four of those features here: region-dependent cell death, time-dependent cell death, T6SS-mediated cell death, and sectoring.

**Figure 3:**
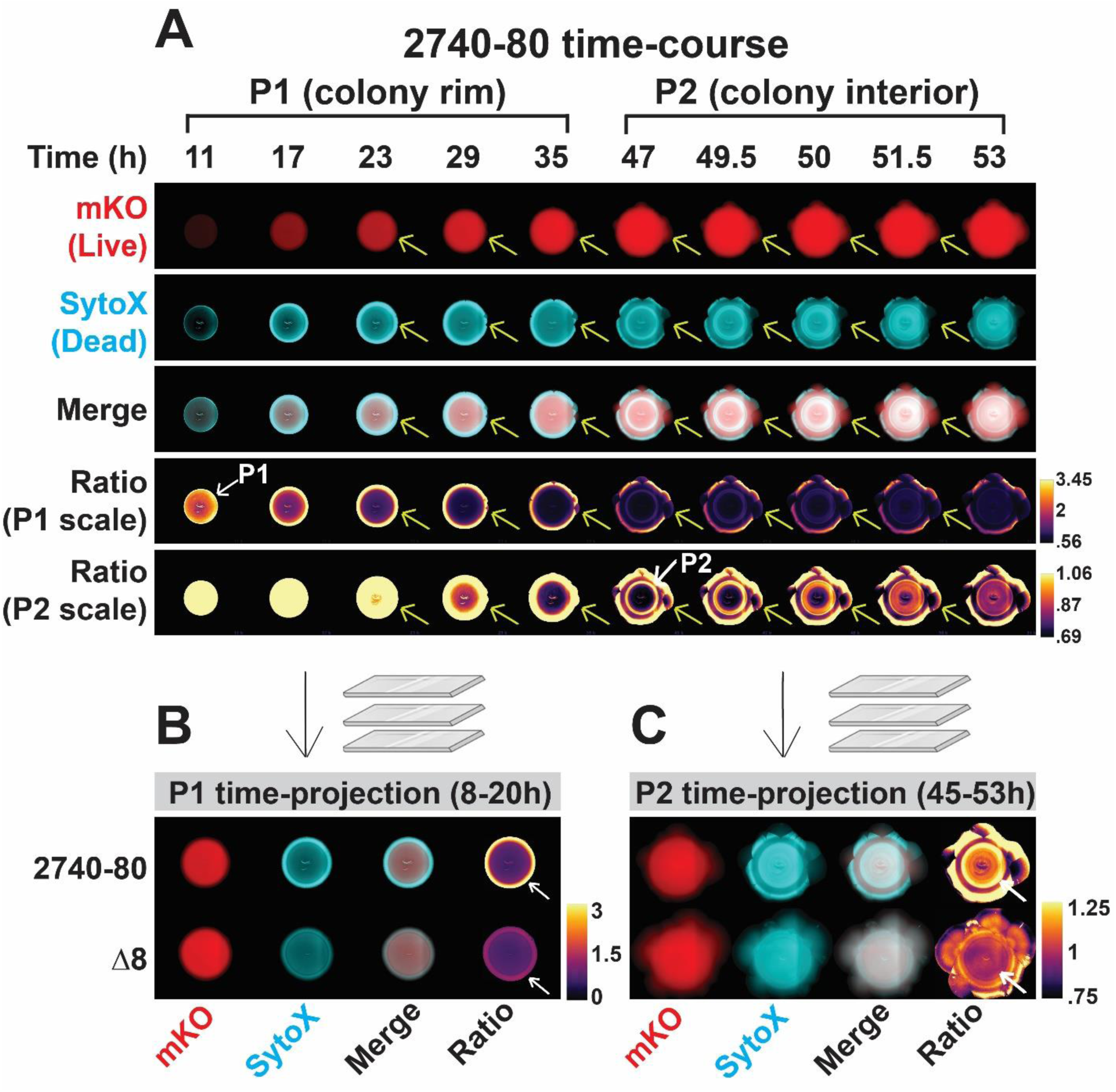
Two phases of spatio-temporal cell death occur in *V. cholerae* 2740-80 and Phase 1 precedes colony sectoring. (A). Images from selected time-points during growth of *V. cholerae* 2740-80 constitutively producing mKO which marks live cells. Dead cells are marked with SytoX stain. Yellow arrows follow one sector as it emerges and expands. Two sets of ratio images are presented to aid visualization of cell death during the Phase 1 (denoted P1 scale) and Phase 2 (denoted P2 scale) time periods. The white arrow with the P1 designation that points to the colony rim highlights the region where maximal Phase 1 cell death occurs. The white arrow with the P2 designation that points to the ring in the colony interior shows the Phase 2 cell death region. In addition to showing each phase of cell death, the P1 scale best depicts that the colony rim is enriched in dead cells while the P2 scale best depicts that the sectors contain primarily live cells. (B,C). Time-projections for Phase 1 (B) and Phase 2 (C) cell death for the indicated strains. White arrows as in A. See also Supplementary Video 1.

**Figure 4.**
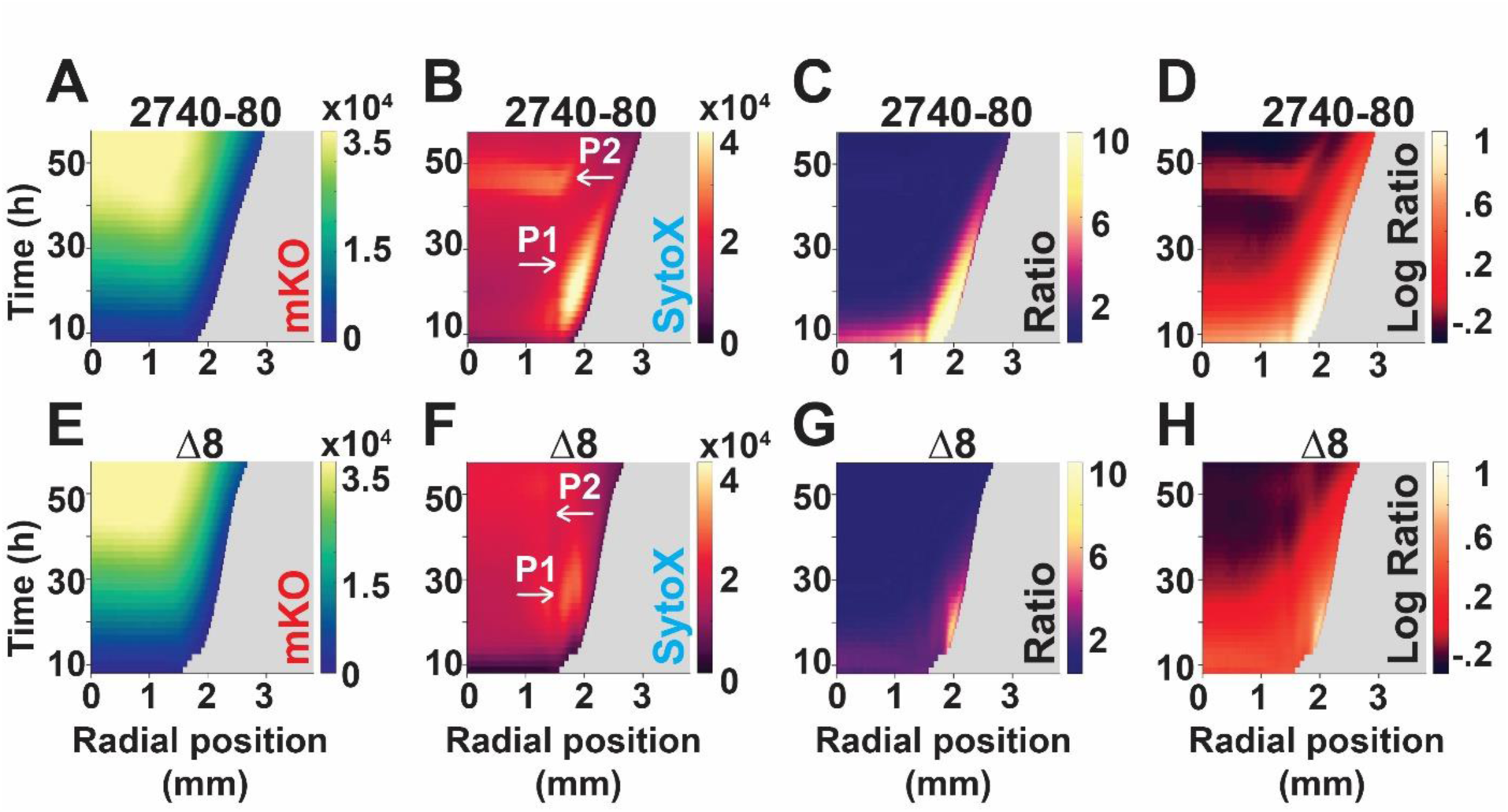
T6SS activity drives each cell death phase in *V. cholerae* 2740-80. (A-H) Space-time kymographs from the indicated channels and strains taken from regions lacking sectors. Kymographs in panels C and G display linear ratio data for visualization of Phase 1 cell death. Logarithmic ratio data are presented in panels D and H for emphasis of Phase 2 cell death. The X-axis on each kymograph indicates the radial position in the colony at which intensity was quantified. The center of the colony is at 0 mm and the colony rim is at ∼3 mm. Phase 1 cell death is indicated with the white arrows labeled P1 in panels B and F and is also visible in panels C,D,G,H along the colony rim as the yellow colored regions. Phase 2 cell death is indicated with the white arrows labeled P2 in panels B and F and is also visible in panel D as the red colored region in the colony interior. In all panels, intensities or ratios are color-mapped and the scale bars represent color:intensity. Intensity ratios were obtained by dividing the intensities from the dead-cell channel by that from the corresponding live-cell channel. Kymographs from one colony are representative of results from 3-9 colonies for each strain.

We first discuss the results from *V. cholerae* 2740-80. The colonies displayed two phases of cell death, which we call Phase 1 and Phase 2, visible in the time-projections and kymographs as regions exhibiting increased SytoX-dependent fluorescence relative to mKO fluorescence. Cell death during Phase 1 occurred between ∼8-40 h and was concentrated predominantly along the periphery of the colony (first four rows in Figure 3A, top row Figure 3B, and Figure 4A-C; indicated with the white arrows and the designation P1; Supplementary Video 1). At ∼44 h, Phase 2 of cell death initiated in the colony interior as a ring and propagated in both the inward and outward directions in an apparent wave-like manner (first three and fifth row in Figure 3A, top row Figure 3C, and Figure 4A,B,D; indicated with the white arrows and the designation P2; Supplementary Video 1). Importantly, ratio-metric kymograph and time-projection analyses of the dead and live cell distributions in the colony (SytoX/mKO) confirmed that the cell death patterns in the different regions and at the different times are not due to differences in cell numbers but, rather, are a consequence of alterations in the ratios of live and dead cells (Figure 3A bottom two rows and Figures 3B,C, 4C,D). The ratio-metric data show that 10-fold more cell death occurs in Phase 1 than in Phase 2, hence, logarithmic ratios of the intensities are provided in Figure 4D to highlight Phase 2 cell death. In all remaining ratio kymographs, we present the log-transformed data. The companion linear ratio data are provided in the Supplemental Information.

Phase 1 cell death largely preceded the formation of sectors, which began along the colony rim (Figure 3A, denoted by yellow arrows). The finding that sector initiation sites co-localize with regions of high Phase 1 cell death is notable given that variants could have emerged anywhere in the colony, as has been observed previously, for example, in *Bacillus subtilis* (Hashuel and Ben-Yehuda, 2019). Rather, in *V. cholerae* 2740-80, sectors arise exclusively in regions of high Phase 1 cell death. Furthermore, cells in the sectors were largely living compared to cells in neighboring non-sectored, parental regions of the colony that were undergoing high cell death (Figure 3A; Supplementary Video 1). This result suggests that the mutations in the arising variant strains suppress the cell death mechanism.

Cell death dynamics were strikingly altered in the T6SS inactive Δ8 strain. Compared to *V. cholerae* 2740-80, Phase 1 cell death along the colony rim was ∼2-10-fold lower in the Δ8 strain and Phase 2 cell death in the colony interior did not occur (Figures 3B,C, 4E-H). Despite displaying decreased cell death, the Δ8 strain developed sectors (Figure 3C). We conclude that the T6SS is involved in driving both phases of cell death in *V. cholerae* colonies. Because sectoring was not abolished in the Δ8 strain, mechanism(s) in addition to T6SS can drive sectoring.

### Distinct T6SS effector-immunity protein pairs drive each phase of *V. cholerae* 2740-80 cell death

We wondered which of the four T6SS effector-immunity (hereafter E-I) protein pairs causes cell death in *V. cholerae* 2740-80. To identify the pair, we engineered strains lacking one (Δ2), two (Δ4), or each combination of three (Δ6) E-I protein pairs. We monitored cell death in the four Δ2 strains, two Δ4 strains, and four Δ6 strains using time-lapse microscopy, as in Figure 3. Data for select strains are displayed in Figure 5. Data for the full set i.e., for each Δ2, Δ4, and Δ6 strain, are displayed in Supplementary Figure 2 (linear ratio kymographs), Supplementary Figure 3 (logarithmic ratio kymographs) and Supplementary Figure 4 (time-series projections).

**Figure 5:**
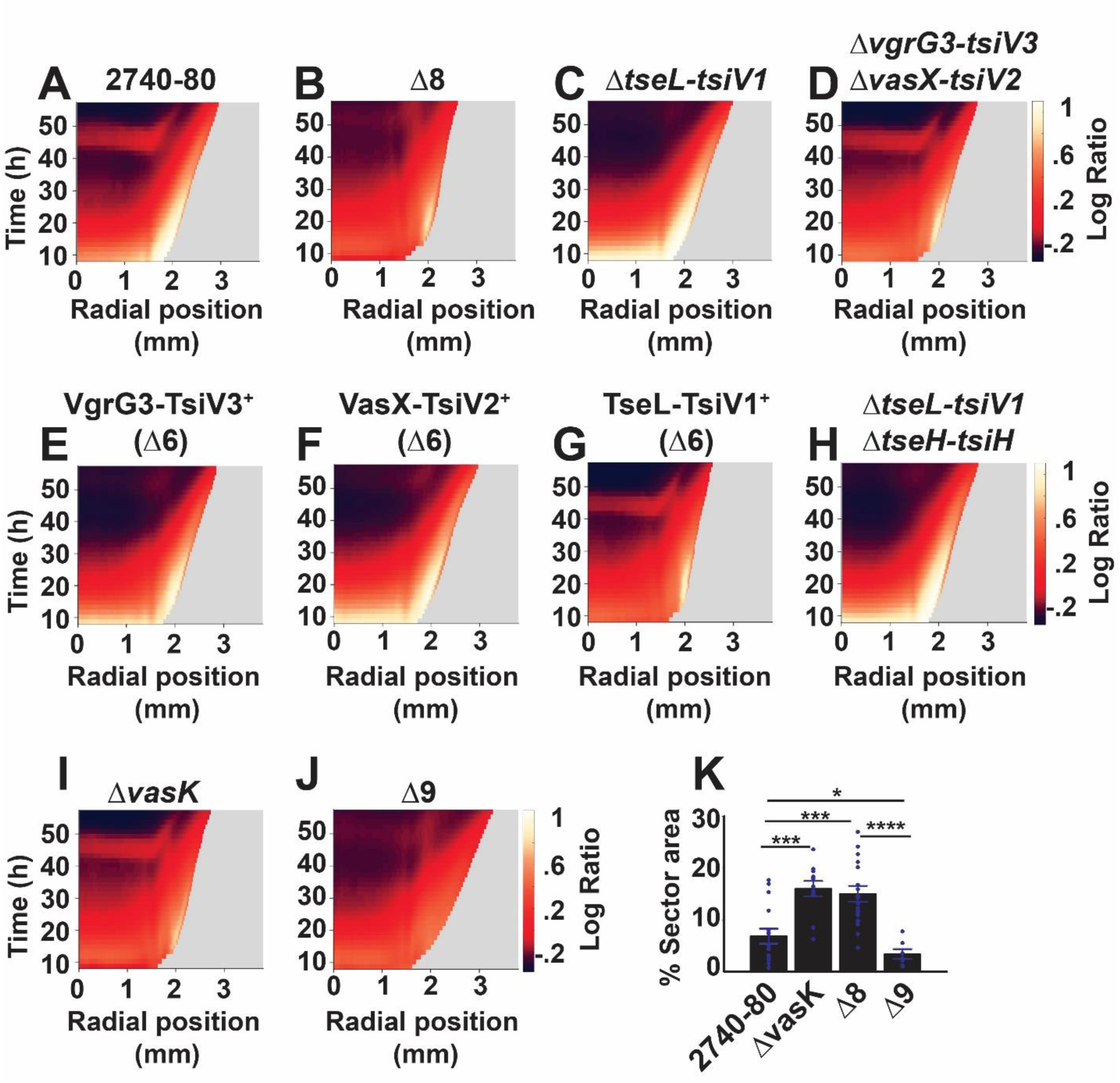
The T6SS apparatus mediates *V. cholerae* colony sectoring, distinct T6SS effector-immunity protein pairs drive each phase of cell death, and Phase 2 cell death does not require the T6SS injection machinery. (A-J) Logarithmic ratio kymographs for the indicated strains. Kymograph data treated as described for Figure 3. Companion linear ratio kymographs for panels A-H are provided in Supplementary Figure 2 and those for panels I-J are in Supplementary Figure 5. (K) Colony area occupied by sectors in the indicated strains. Kymographs from one colony are representative of results from 3-9 colonies for each strain. In panel K, data represent average values from biological replicates (n=7-16), and error bars show SEMs. Statistical significance was calculated using a two-tailed Student’s *t* test with unequal sample variance. Asterisks are as follows: * *P* < 0.05, **** *P* < 0.005, **** *P* < 0.0005.

Each Δ2 strain, lacking one E-I protein pair, displayed Phase 1 cell death along the colony rim that was indistinguishable from that of *V. cholerae* 2740-80 (Supplementary Figures 2A-F, 3A-F and Supplementary Figure 4A). Data from the strain lacking the TseL-TsiV1 pair is provided as the representative of the Δ2 strains in Figure 5C and should be compared to the data in Figure 5A,B for *V. cholerae* 2740-80 and the Δ8 strain, respectively. The Δ4 strain lacking both the VgrG3-TsiV3 and the VasX-TsiV2 protein pairs, by contrast, showed the reduced Phase 1 cell death phenotype of the Δ8 strain (Figure 5B,D and panels B,G in Supplementary Figures 2, 3, and Supplementary Figure 4A). Because the Δ2 strains did not display defects in Phase 1 cell death, while the Δ4 strain was impaired for killing, we conclude that the VgrG3 and VasX effector proteins make redundant contributions to Phase 1 cell death. Indeed, confirming this assertion, among the Δ6 strains, only two strains, harboring either VgG3-TsiV3 or VasX-TsiV2 as the sole E-I protein pair, displayed Phase 1 cell death patterns like *V. cholerae* 2740-80 (Figure 5A,E,F,G and panels A,H,I,J,K in Supplementary Figures 2, 3, and Supplementary Figure 4A). Consistent with the idea that the remaining two E-I protein pairs, TseL-TsiV1 and TseH-TsiH, are dispensable for driving Phase 1 cell death, the Δ4 strain lacking both of these protein pairs acted like *V. cholerae* 2740-80 with respect to Phase 1 cell death (Figure 5A,H and panels A,L in Supplementary Figures 2, 3, and Supplementary Figure 4A). Thus, either the VgG3 or the VasX effector can mediate Phase 1 cell death.

Regarding Phase 2 cell death, which occurs in the colony interior, among the Δ2 strains, cell death was abolished only in the strain lacking TseL-TsiV1, a phenotype mimicking the Δ8 strain (compare data in Figure 5C in the colony interior to that in panels 5A and 5B and see panels A-F in Supplementary Figures 2, 3, and Supplementary Figure 4B). Further confirming the role of the TseL-TsiV1 protein pair in Phase 2 cell death, only one Δ6 strain, possessing TseL-TsiV1 as the sole E-I protein pair, showed a Phase 2 cell death pattern akin to that of *V. cholerae* 2740-80 (Figure 5A,E,F,G and panels A,H,I,J,K in Supplementary Figures 2, 3, and Supplementary Figure 4B). Thus, the TseL effector protein is required to drive Phase 2 cell death.

### Phase 2 cell death does not require the T6SS injection machinery

T6SS-dependent killing relies on an injection machine to deliver toxic effector proteins into prey cells. To examine whether the T6SS-dependent killing that takes place in the *V. cholerae* 2740-80 colonies requires the T6SS-injection apparatus, we deleted *vasK*, encoding an essential structural component of the injection machine from *V. cholerae* 2740-80 and from the Δ8 strain. Phase 1 killing along the colony rim was diminished in both the Δ*vasK* and the Δ8 strains and combining the Δ*vasK* and Δ8 mutations (hereafter the Δ9 strain), reinforced the other’s effects, nearly eliminating Phase 1 cell death (Figure 5A,B,I,J; Supplementary Figures 5A-D and Supplementary Figure 6A). Thus, possession of a functional T6SS-injection machinery contributes strongly to Phase 1 cell death. We offer possibilities that could account for the synergistic effects of the combined Δ*vasK* and Δ8 mutations in the discussion.

With respect to Phase 2 cell death in the colony interior, the Δ*vasK* strain displayed no defect while Phase 2 cell death did not occur in the Δ8 and Δ9 strains (Figure 5A,B,I,J; Supplementary Figure 6B). Since Phase 2 cell death is driven by the TseL effector protein (Figure 5C,G,H), we conclude that TseL can cause cell death independent of the T6SS-injection apparatus. This experiment does not allow us to distinguish between whether TseL is translocated to target cells via an alternate mechanism or whether TseL-producing cells experience auto-poisoning.

### T6SS-activity drives colony sectoring

To probe whether T6SS activity influences sectoring, we assayed the phenotypes in the strains under study. *V. cholerae* 2740-80, strains lacking individual or combinations of E-I protein pairs, the Δ8 strain, and the Δ*vasK* strain all made sectors (Supplementary Figure 4B and Supplementary Figure 6B). By contrast, the Δ9 strain consistently formed fewer and/or smaller sectors (Supplementary Figure 6B). Using machine-learning-driven image segmentation, we measured the area occupied by sectors in the Δ9 strain and its progenitors. Sectors in the Δ*vasK* and Δ8 mutants occupied ∼2.5-fold more area than did sectors in *V. cholerae* 2740-80 (Figure 5K). Sectors in the Δ9 mutant occupied ∼2-fold less area than sectors in *V. cholerae* 2740-80 and ∼5-fold less area than in the Δ*vasK* and Δ8 strains (Figure 5K). We conclude that T6SS-killing activity drives colony sectoring. In the Discussion, we present possible explanations for the unexpected finding that the Δ*vasK* and Δ8 mutations each drive increases in sector area occupancy while, when combined, they reduce the colony area occupied by sectors.

### The LCD QS state eliminates colony sectoring through repression of T6SS-dependent cell killing and activation of Vps-dependent T6SS defense

The above results suggest that T6SS plays a key role in causing cell death and colony sectoring. As mentioned, excess Vps can defend against T6SS killing. We know that QS controls both *t6ss* and *vps* expression in *V. cholerae*. This understanding enables us to put forward and test the idea that the QS LCD-locked variants we isolated exhibit both reduced T6SS activity and high Vps production. Together, these altered traits suppress T6SS-dependent killing in the colony, which decreases overall cell death, the consequence of which is prevention of sectoring.

To assess whether the LCD QS state alters *V. cholerae* 2740-80 T6SS activity, we measured the capacity of the *luxO* variants we isolated to kill *Escherichia coli* in an inter-bacterial T6SS-dependent killing assay. As prey, we used an *E. coli* strain that constitutively produces luciferase (*lux*), and thus, light output tracks with live prey cells. When the *luxO* variants were used as predators, there was a 10-100-fold decrease in prey killing relative to when *V. cholerae* 2740-80 was predator (Figure 6A). No killing occurred when the Δ*vasK* strain was the predator, confirming that killing requires T6SS activity (Figure 6A). To verify that the decreases in killing ability of the *luxO* variants were a consequence of decreased expression of *t6ss* genes, in one representative *luxO* variant (*luxO* A97E), we quantified transcript levels for the genes specifying each E-I protein pair and select genes encoding T6SS structural components. Figure 6B shows the results. Compared to *V. cholerae* 2740-80, the *luxO* A97E variant exhibited 2-4-fold decreased expression of every tested *t6ss* gene. Thus, LuxO-driven LCD behavior suppresses T6SS-killing activity in *V. cholerae* 2740-80.

**Figure 6:**
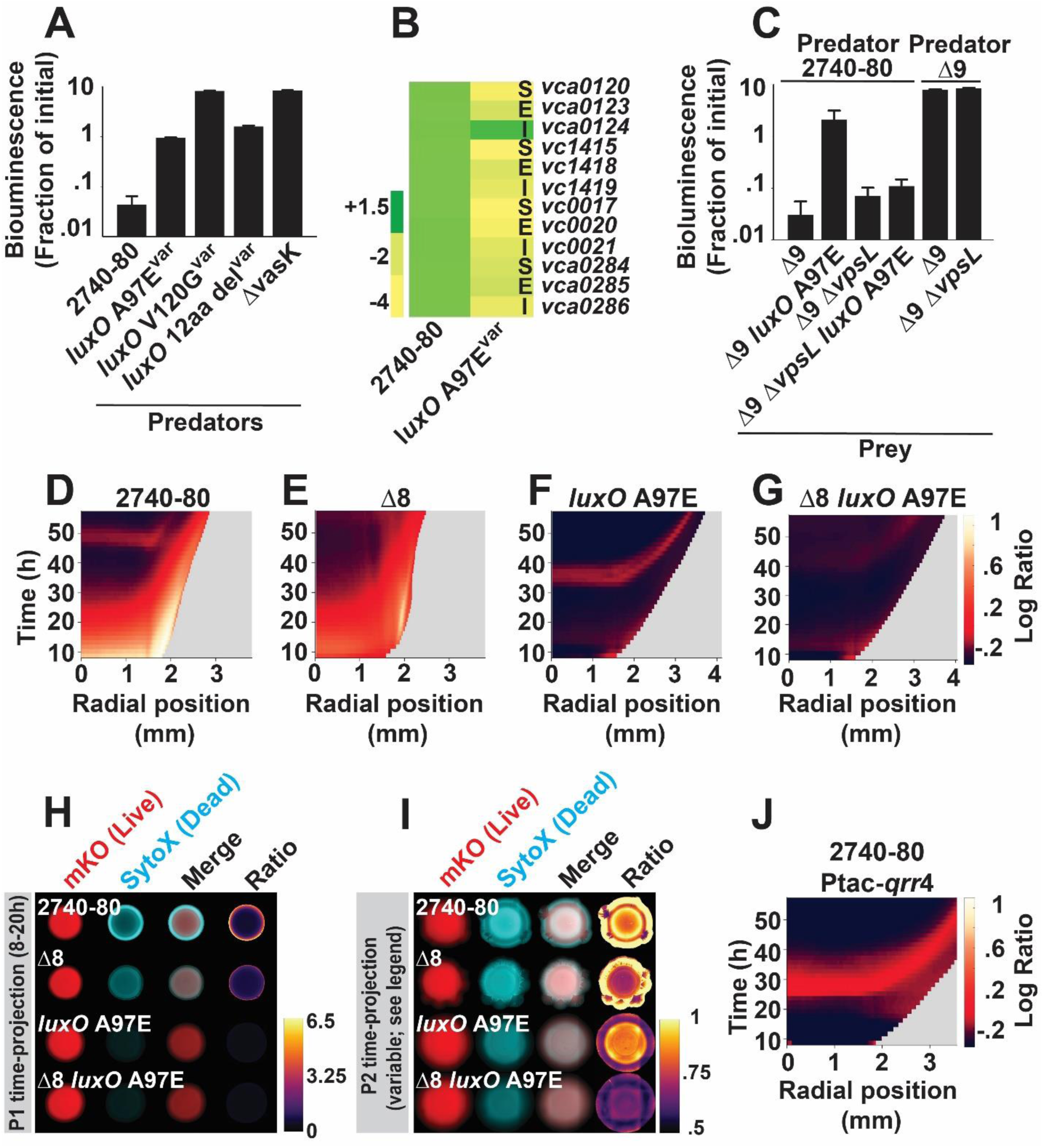
The LCD QS state represses T6SS-dependent killing, activates Vps-dependent T6SS-defense, and eliminates colony sectoring. (A). Inter-bacterial T6SS killing assay measuring survival of T6SS-inactive *E. coli* prey following challenge with the indicated *V. cholerae* predator cells. The *E. coli* prey strain constitutively expresses luciferase. Thus, bioluminescence output is a proxy for live cells. (B) Transcript abundance, relative to *V. cholerae* 2740-80, for the indicated strains and genes. The data are color-mapped, and the scale bar displays fold-change. Designations: S=T6SS secretion protein, E=T6SS effector toxin, I= T6SS immunity protein. (C) As in A, measuring survival of the indicated *V. cholerae* prey strains following challenge with the designated *V. cholerae* 2740-80 predator strains. (D-G) Logarithmic ratio kymographs for the indicated strains. (H,I) Time-projections showing cell death and sectoring for the indicated strains and phases. (J) Logarithmic ratio kymograph for *V. cholerae* 2740-80 in which *qrr*4 is overexpressed. Kymograph data in panels D-G and J treated as described for Figure 3. Kymographs from one colony are representative of results from 3-9 colonies for each strain. In A and C, data represent average values from biological replicates (n=3), and error bars show SDs. In B, average values were obtained from three biological replicates and two technical replicates for each strain (n=6). In I, due to differences in timing of Phase 2 cell death among strains (compare Phase 2 initiation times in panels D and F), time-projections for *V. cholerae* 2740-80 and the Δ8 strain show data from 42.5-56 h, while for the *luxO* A97E and Δ8 *luxO* A97E strains, data are shown from 30.5-44 h.

We examined whether increased Vps production boosts the defense capacity of the *V. cholerae* 2740-80 LCD-locked QS variants against incoming T6SS attacks. We already know that all the *V. cholerae* 2740-80 LCD-locked QS variants exhibit increased expression of *vps* (Figure 2D), so we again used the *luxO* A97E allele as our representative for this analysis. To do the experiment, we introduced alone and in combination, the *luxO* A97E and Δ*vpsL* mutations into the Δ9 strain. This strategy allowed us to avoid possible complications from secondary mutations that might be present in the original *luxO* A97E variant strain. Moreover, because each strain in this set lacks all T6SS-immunity proteins, they are susceptible to T6SS-dependent killing following challenge with *V. cholerae* 2740-80. Lastly, all prey strains were also engineered to carry a constitutive *lux* reporter enabling tracking of survival. Relative to the Δ9 strain, the Δ9 *luxO* A97E strain displayed ∼100-fold increased survival against the *V. cholerae* 2740-80 predator while the Δ9 Δ*vpsL* and Δ9 Δ*vpsL luxO* A97E strains showed no survival enhancement (Figure 6C). No killing occurred when the Δ9 or Δ9 Δ*vpsL* strains were challenged with the Δ9 strain as predator, again confirming that T6SS activity drives killing in our assay (Figure 6C). Thus, in the *luxO* A97E LCD-locked variant, and presumably the other variants we isolated, increased Vps production driven by QS functioning in the LCD-mode drives enhanced defense against incoming T6SS attacks. Moreover, the results show that the protective effect of high level Vps production can overcome the sensitivity to killing caused by complete lack of immunity factors.

Beyond effects on T6SS-killing and T6SS-defense, we examined whether the LCD-locked QS mode promotes an altered cell death pattern. Here, we again use the *luxO* A97E mutation as the representative, and as above, to avoid complications from possible secondary mutations in the original variant, we reconstructed all needed mutations in *V. cholerae* 2740-80 and in the Δ8 strain. Phase 1 cell death along the colony rim was abolished in the *luxO* A97E and Δ8 *luxO* A97E strains (Figure 6 D-H and Supplementary Figure 5A,B,E,F). Indeed, the *luxO* A97E mutation eliminated the residual Phase 1 cell death that occurs in the Δ8 strain (Figure 6E,F,G). Thus, QS fully controls Phase 1 cell death. Given that elimination of T6SS does not abolish all Phase 1 cell death in the Δ8 strain (Figure 6E), another QS-controlled process must be involved in Phase 1 cell death. By contrast, the *luxO* A97E strain displayed Phase 2 cell death in the colony interior like *V. cholerae* 2740-80 (Figure 6 D,F,I). The Δ8 *luxO* A97E strain mimicked the Δ8 strain and displayed no Phase 2 killing (Figure 6E,F,G,I). Thus, Phase 2 cell death, which is TseL-dependent (Figure 5C,G,H), is not subject to QS regulation. Finally, none of the strains carrying *luxO* A97E sectored, including in the Δ8 background (Figure 6I; see bottom two rows). Thus, the LCD QS state is epistatic to T6SS with respect to sectoring. Since the LCD-locked variants do not sector and have no Phase 1 cell death, but undergo Phase 2 cell death, we infer that Phase 1 cell death is key to the sectoring phenotype while Phase 2 cell death may be dispensable for sectoring.

To pinpoint the mechanism that connects QS to T6SS-driven cell death, Vps, and sectoring phenotypes, we focused on the Qrr small RNAs that repress translation of the large *t6ss* gene cluster, and indirectly activates *vps* gene expression (Figure 1) (Shao and Bassler, 2014). To test if the QS phenotypes hinge on Qrr activity, we constitutively expressed one of them, *qrr*4 (Ptac-*qrr*4), in *V. cholerae* 2740-80 and examined the phenotypic consequences. Introduction of Ptac-*qrr*4 abolished Phase 1 cell death and sectoring in *V. cholerae* 2740-80 including the low-level cell death that occurs in the Δ8 strain (Figure 6J and Supplementary Figure 5G). Phase 2 cell death occurred (Figure 6J). Thus, overexpression of Qrr4 is sufficient to mimic the phenotype caused by the *luxO* A97E mutation (Figures 6F,J). We conclude that in our LCD-locked *luxO* QS variants, it is the Qrr sRNAs that repress T6SS components and activate Vps production. Together, these changes lower cell death and abolish sectoring.

### Constitutive expression of *t6ss* genes eliminates the spatio-temporal cell death patterns in *V. cholerae* 2740-80 and restores cell death in the LCD-locked QS strain

Our data suggest that QS-dependent spatio-temporal control of T6SS activity and Vps production influences cell death patterning and sectoring in *V. cholerae* 2740-80. If so, we can make two predictions: First, forcing production of T6SS machinery in all cells in *V. cholerae* 2740-80 colonies would cause cell death across the entire population and eliminate any spatio-temporal pattern. Second, re-establishment of T6SS production in a LCD-locked QS strain would restore cell death and drive sectoring. To test the first prediction, we introduced a plasmid carrying the T6SS activators *qstR* and *tfoX* under control of an arabinose inducible promoter (called P*t6ss*-ON) into *V. cholerae* 2740-80 (Bernardy et al., 2016; Jaskólska et al., 2018; Metzger et al., 2016). To test the second prediction, we did the same experiment in the LCD-locked *luxO* A97E strain. In each case, we monitored cell death and sectoring.

Induction of P*t6ss*-ON-driven T6SS production caused a 10-40-fold increase in cell death in *V. cholerae* 2740-80 compared to the strain carrying the empty vector (Figure 7A-D). Notably, *V. cholerae* 2740-80 harboring the empty vector displayed the characteristic Phase 1 and Phase 2 cell death patterns, while introduction of P*t6ss*-ON caused cell death across the colony (Figure 7C,D). Thus, the normal patten of cell death that occurs in *V. cholerae* 2740-80 colonies is a consequence of non-homogeneous *t6ss* expression and the ensuing non-homogeneous T6SS activity.

**Figure 7:**
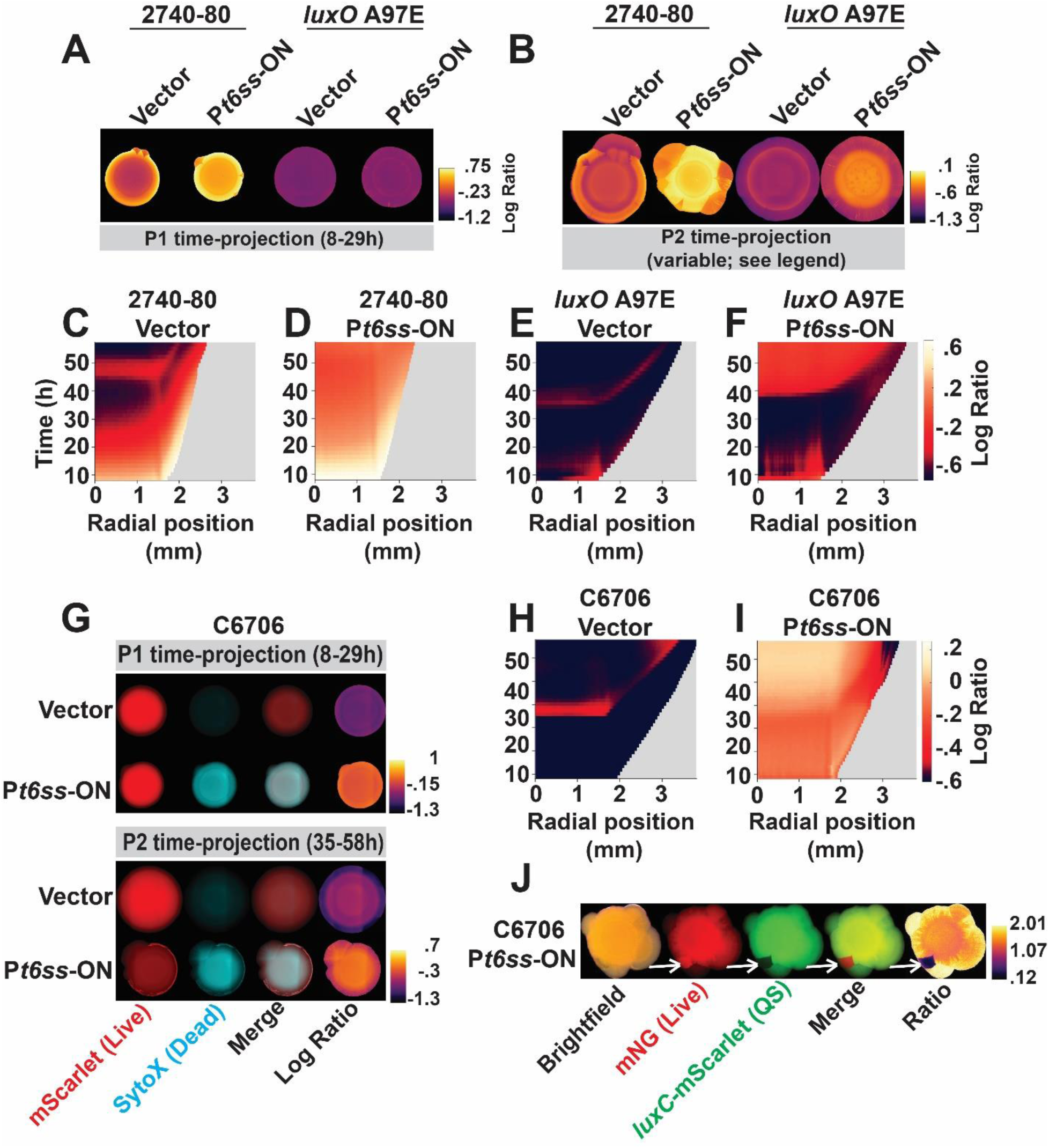
Inducible *t6ss* expression eliminates the cell death pattern in *V. cholerae* 2740-80 and drives sectoring and the emergence of QS variants in the normally T6SS-silent *V. cholerae* C6706 strain. (A,B) Logarithmic ratio time-projections showing cell death and sectoring for the indicated phases in strains carrying either an empty vector (denoted: Vector) or the vector carrying P*t6ss*-ON. Phase 2 projection timing as in Figure 6I. (C-F) Logarithmic ratio kymographs for the indicated strains carrying the designated plasmids. (G) Time-projections showing cell death and sectoring in *V. cholerae* C6706 carrying the indicated plasmids (H,I) Logarithmic ratio kymographs for the indicated strains. (J) Stereo-microscope images of *V. cholerae* C6706 carrying P*t6ss*-ON, constitutively producing mNeonGreen (denoted: mNG) to mark live cells, and a QS-activated reporter (*luxC*-mScarlet). White arrows mark a sector that contains live cells but exhibits little *luxC*-mScarlet activity. Kymograph data in panels C-F and H,I treated as described for Figure 3. Kymographs from one colony are representative of results from 3-9 colonies for each strain. Note: because plasmid-driven production of T6SS components drives large increases in cell death, the ratio time-projection images in A, B, G are displayed on log scales, unlike those in Figures 3, 5, and 6 which are shown on linear scales.

Introduction of P*t6ss*-ON into the *luxO* A97E strain caused little cell death until ∼40 h, after which cell death became detectable and occurred homogenously across the colony interior. The colony rim did not display cell death (Figure 7A,B, E,F). Once cell death commenced, the level was roughly the same as that in *V. cholerae* 2740-80 carrying P*t6ss*-ON. Because Qrr4 is constitutively produced in the *luxO* A97E LCD-locked QS strain and because P*t6ss*-ON activates transcription of *t6ss* genes, the absence of cell death during the 8-40 h timeframe and along the colony rim suggests that, during early times, *t6ss* genes remains subject to Qrr4-mediated post-transcriptional repression. During later growth times in the colony interior, some process must override Qrr4 regulation and allow P*t6ss*-ON-dependent T6SS-activation to restore cell death in the LCD-locked strain.

Regarding sectoring, *V. cholerae* 2740-80 carrying P*t6ss*-ON formed sectors, whereas only minimal sectoring occurred following introduction of P*t6ss*-ON into the *luxO* A97E strain, visible as radial streaks (Figure 7B and see also enlarged images in Supplementary Figure 6C). We do not understand why sectoring was not fully restored. Likely, plasmid expression of the genes encoding the two T6SS activators does not perfectly mimic native control of the entire set of *t6ss* gene clusters. Nonetheless, we conclude that QS governs the region-specific expression of Phase 1 T6SS activity thereby driving cell death and sectoring.

### T6SS-dependent cell death, sectoring, and emergence of QS variants occurs when *t6ss* genes are expressed in a normally T6SS-silent *V. cholerae* strain

To garner additional evidence demonstrating that both cell death and sectoring are T6SS-dependent in *V. cholerae*, we used our P*t6ss*-ON construct to induce *t6ss* gene expression in *V. cholerae* C6706, which as mentioned, does not express *t6ss* genes under laboratory growth conditions and does not sector (Figure 2A). We assessed the consequences to cell death and sectoring. *V. cholerae* C6706 carrying the empty vector displayed no Phase 1 cell death (Figure 7G top row and panel H). There was modest Phase 2 cell death, but notably, ∼10-fold lower than that in *V. cholerae* 2740-80 (compare data in Figure 7G third row and panel H to that in Figure 7B,C). Figure 7G,H,I show that P*t6ss*-ON-driven T6SS production increased cell death ∼10-fold in *V. cholerae* C6706. Cell death occurred across the entire colony, consistent with homogeneous expression of *t6ss* genes. Furthermore, sectors formed with timing similar to that in *V. cholerae* 2740-80 (Figure 7G, bottom row). Thus, T6SS activity causes cell death and the appearance of sectors in both pandemic (*V. cholerae* C6706) and pre-pandemic (*V. cholerae* 2740-80) *V. cholerae* strains.

To discover whether the *V. cholerae* C6706 T6SS-dependent sectors are enriched in cells with altered QS behaviors, we imaged sectors in a *V. cholerae* C6706 strain harboring P*t6ss*-ON, a constitutively produced fluorescent reporter marking live cells (mNG), and a QS-activated-fluorescent reporter (*luxC*-mScarlet). In ∼5-10% of the sectors, the live cells present did not express the QS reporter, indicating that the cells in these sectors had acquired mutation(s) that result in LCD-locked QS behavior (Figure 7J; indicated with white arrows). We conclude that T6SS killing activity in *V. cholerae* colonies imposes a selective pressure to acquire LCD locked QS mutations, presumably enhancing growth and promoting sector formation.

## Discussion

Here, we discover that QS-controlled T6SS-mediated cell death provides a selective pressure that allows QS defective strains of *V. cholerae* to arise that are capable of evading T6SS-killing. T6SS-mediated cell death occurs in a two-phase, spatio-temporal manner. Distinct T6SS effectors, VgrG3 and VasX for Phase 1 and TseL for Phase 2, are required for killing. QS controls Phase 1 cell death. Phase 1 cell death is key for sectoring to occur and thus, for enhanced genetic diversity to arise in the population (see model in Supplementary Figure 7).

Our findings reveal an unanticipated new facet of *V. cholerae* T6SS biology: the *V. cholerae* T6SS machinery, which was understood to deliver toxins to non-kin cells, can be deployed to eliminate sibling cells. Thus, the T6SS machinery may have unappreciated roles in intra-specific antagonism. It was surprising that sibling cells succumb to incoming T6SS attacks given that they produce T6SS-effector neutralizing immunity proteins. One previous example of T6SS-dependent kin-killing has been reported in *Myxococcus xanthus*, in which slow-growing or auxotrophic cells in the population exhibit reduced T6SS protein production, including T6SS immunity proteins, rendering them susceptible to killing by faster growing nearby cells that produce higher levels of T6SS toxins (Troselj et al., 2018). Because Phase 1 cell death in *V. cholerae* colonies occurred primarily along the rim of the colonies i.e., in the region experiencing maximal growth, differences in growth seem an unlikely explanation for the alterations in *t6ss* gene expression patterns we observe, contrasting our findings from those in *M. xanthus*. Rather, we propose that, during the Phase 1 cell death period, heterogeneous *t6ss* gene expression occurs in *V. cholerae* colonies due to gradients of QS autoinducers, although we caution that this hypothesis remains to be tested. In support of this argument, introduction of the P*t6ss*-ON plasmid into wildtype *V. cholerae* induced homogeneous cell death across the colony, including in regions of the colony that do not normally undergo cell death (Figure 7D). Likewise, in the locked LCD *V. cholerae luxO* A97E strain, which has overall reduced T6SS activity and displays no detectable Phase 1 cell death, introduction of P*t6ss*-ON also elicited homogeneous killing (Figure7F). Together, these results suggest that the wildtype killing pattern stems from distinct levels of expression of *t6ss* genes in restricted regions of the colony. An alternative explanation is that a sub-population of *V. cholerae* cells does not properly express genes required for *t6ss* defense, such as those involved in production of Vps or that defend against T6SS-imposed membrane stress (Figure 6C and (Hersch et al., 2020)).

*V. cholerae* colonizes chitinous surfaces in its marine environment and chitin acts as a cue that activates *t6ss* gene expression (Borgeaud et al., 2015; Meibom et al., 2005). Curiously, environmental isolates of *V. cholerae* often harbor QS inactivating mutations (Joelsson et al., 2006). Our results demonstrate that, under laboratory growth conditions, T6SS-driven killing fosters the emergence of QS LCD-locked variants in *V. cholerae* colonies. We propose that T6SS-driven kin-killing likely also occurs in natural habitats and perhaps during disease, and this mechanism propels genetic diversity. It is also intriguing that the arising variant strains exhibit a range of T6SS activity levels and/or capacities to neutralize incoming T6SS attacks (Figure 6A,C). Such a possibility also exists for the other two variants we recovered in our suppressor screen, which had acquired mutations affecting CspA and PyrG function. CspA, a cold shock protein, was previously shown to modulate T6SS killing activity, while the PyrG cytidine synthase likely affects T6SS function by altering levels of cytidine, a ligand for the CytR transcription factor, the function of which is to activate *t6ss* genes while repressing genes required for biofilm formation (Barbier et al., 1997; Townsley et al., 2016; Watve et al., 2015). Thus, T6SS-driven intra-specific antagonism selects for acquisition of mutations in QS and other pathways that modify expression of *t6ss* offensive and defensive genes. This mechanism could enable iterative improvements in the ability of *V. cholerae* to tune T6SS activity to its various niches.

We discovered that T6SS-driven Phase 1 cell killing relies on the T6SS VasK-dependent injection machinery (Figure 5I). Curiously, combining the Δ8 mutation, which eliminates all effectors, with the Δ*vasK* mutation, eliminating the injection apparatus, had a modest additive effect with respect to cell death (additivity is best visualized in Supplementary Figure 5B-D). Two possible explanations occur to us. First, despite lacking T6SS toxins, the Δ8 strain nonetheless possesses an intact T6SS injection machine. It is possible that a subset of cells in the colony have damaged cell envelopes, rendering them susceptible to harm upon physical penetration by the T6SS needle, which is expelled with considerable energy into target cells (Kamal et al., 2020; Wang et al., 2019). Emphasizing this line of thought, a recent study found that *V. cholerae* cells possessing only the injection machinery, but no effectors, can inhibit the growth of *P. aeruginosa* strains lacking the TolB protein, which is important for maintaining outer membrane integrity (Kamal et al., 2020). A second possibility is that the *V. cholerae* Δ8 strain continues to synthesize an as yet unidentified effector toxin that employs the T6SS injection apparatus for its killing activity.

Phase 2 cell killing required the T6SS TseL effector toxin but not the T6SS injection machinery (Figure 5C,I). It is currently unclear whether TseL causes self-killing or if it can be secreted via an alternate secretion mechanism. In support of the notion that TseL contributes to self-killing, Ho et al. (2017) showed that TseL can be trafficked from the cytosol to the periplasm via a non-T6SS-dependent mechanism. Thus, one possibility is that time- and region-specific trafficking of TseL to the periplasmic compartment promotes Phase 2 cell death. TseL is a phospholipase (Dong et al., 2013). An alternative possibility is that TseL residing in the cytoplasm destroys essential cytoplasmic factor(s), such as precursors in phospholipid biosynthesis, the absence of which would cause cell death.

In multicellular organisms, including humans, key segments of development rely on genetically regulated and time- and region-specific cell death processes (Fink and Cookson, 2005; Kerr et al., 1972). In a striking parallel, we show here that cell death in *V. cholerae* colonies is QS-regulated and occurs in a time- and region-specific manner. Cell death wave(s) were recently reported to guide eukaryotic apoptosis in *Xenopus laevis* (African frog) eggs (Cheng and Ferrell, 2018). Our time-lapse videos and kymograph analyses hint that, in, *V. cholerae*, cell death during Phase 2 may also propagate as a wave (see especially Supplementary Video 1). While currently speculative, if so, this feature would mirror what occurs in eukaryotes. We are currently exploring the origin of the wave-like behavior observed here.

Regarding the sequential timing of the two phases of cell death in *V. cholerae* 2740-80, we note that Phase 2 cell death commences only after Phase 1 death subsides. Also, in QS LCD-locked strains (*luxO* A97E or *V. cholerae* 2740-80 carrying Ptac-*qrr*4), which lack Phase 1 cell death, the timing of onset of Phase 2 cell death shifts dramatically, initiating ∼24 h earlier that in a strain that is wildtype for QS (compare timing in Figure 6D to that in Figure 6J). Thus, it appears that the timing and occurrence of Phase 1 killing sets the timing of Phase 2 killing. It could be that Phase 1 killing, which occurs at the colony rim among the youngest members of the colony, functions to delay cell death in the population elders, as Phase 2 killing occurs in the colony interior which contains the oldest cells in the colony. Possibly, cells undergoing death at the colony rim release a “defer/delay” signal that is detected by cells in the colony interior. Such a signal, by alerting older cells to impending cell death, could function to buy them time to protect themselves. If so, such a scenario would present another fascinating parallel to eukaryotic cell death where, following initiation of apoptosis, dying cells release chemical signals that are detected by stem cells, prompting the stem cells (which are the oldest cells in eukaryotic tissue communities) to mount defenses that ensure their survival and, in turn, their capacity for future tissue re-population (Xing et al., 2015).

## Materials and Methods

### Materials

Kits for gel purification, plasmid-preparation, RNA-preparation (RNeasy), qRT-PCR, and RNA-Protect reagent were purchased from Qiagen. iProof DNA polymerase and deoxynucleoside triphosphates were purchased from Biorad.

### Bacterial growth

*Saccharomyces cerevisiae* and *E. coli* Top10 were used for cloning. *E. coli* S17-1 λ*pir* was used for conjugations. Cultures of *V. cholerae* and *E. coli* were grown in LB medium at 37°C with shaking, with a headspace to growth medium volume ratio of 7. The only exception is that prey strains for killing assays were grown overnight at 30°C. When required, media were supplemented with streptomycin, 200 μg/mL; kanamycin 50 μg/mL; polymyxin B, 50 μg/mL; chloramphenicol, 1 μg/mL; spectinomycin, 200 μg/mL. In experiments requiring induction of gene expression, all media used were supplemented with 0.1% arabinose. All *V. cholerae* assays were performed at 30°C unless otherwise noted. LB medium (both liquid and solid) was prepared with dd H_2_O, 100 % Tap H_2_O, or 80% Tap and 20% dd H_2_O. Changes in media preparation were a consequence of COVID-imposed supply issues and LB reagent acquired from multiple suppliers. Differences in batches affected the timing of assays and amount of sectoring.

Consistent phenotypes could be obtained when solid LB medium was prepared with 80% Tap and 20% dd H_2_O and liquid LB medium was prepared with 100% Tap H_2_O. Bioluminescence-reporter assays were conducted as previously described (Mashruwala and Bassler, 2020).

### Strain construction

Chromosomal alterations in *V. cholerae* strains were introduced using the pRE112 suicide vector harboring the counter-selectable *sacB* gene as previously described (Edwards et al., 1998; Eickhoff et al., 2021). All strains used in the study are listed in Supplementary Table 2. Unless otherwise specified, chromosomal DNA from *V. cholerae* 2740-80 was used as template for PCR reactions. Plasmids were constructed using pBAD-pEVS or pRE112 as backbones and assembled using enzyme-free XthA-dependent *in vivo* recombination cloning or yeast-recombination-assisted assembly as previously described (Beyer et al., 2015; Joska et al., 2014; Mashruwala and Boyd, 2016; Nozaki and Niki, 2018). Plasmids were introduced into *V. cholerae* strains by conjugation with *E. coli* S17-1 λ*pir*, as described previously. Plasmids used in this work are listed in Supplementary Table 3.

### Colony sectoring and cell death assay

A 700 µL aliquot of an overnight culture of *V. cholerae* was combined with 4 mm glass beads in an Eppendorf tube and subjected to vortex for 5 min to disperse aggregates. The culture was diluted with 1X PBS to a final OD_600_ of 0.5. The sample was again subjected to vortex, without glass beads, for 5 min. A 1 µL aliquot of this suspension was spotted onto 35 mL of solid LB agar in a one well plate and allowed to dry for 5 min at room temperature. The plate was incubated for the remainder of the assay at 30°C. Up to 24 such samples were aliquoted onto each agar pad. Sector formation became visible between 18-48 h. When required, the LB agar medium was supplemented with 2 µM SytoX dye (ThermoFisher) (Asally et al., 2012).

### Bioluminescence-based T6SS-dependent interbacterial killing assay

Prey cells constitutively expressed the *luxCDABE* operon, incorporated onto the chromosome (*V. cholerae*) or a plasmid (*E. coli* Top10). Prey cell light production was quantified to track surviving cells. To initiate the killing assay, 800 µL of overnight cultures of prey and predator strains were concentrated two-fold by centrifugation and resuspension in 400 µL 1X PBS. The predator cell suspension was combined with 4 mm glass beads in an Eppendorf tube and subjected to vortex for 5 min to disperse aggregates. In experiments in which effects of Vps production on T6SS-driven killing were examined, rather than apply vortex, cells were gently resuspended with a pipette to preserve biofilm structures. In the case of prey, cultures were divided in half. One half was subjected to vortex, as described above, and used to obtain the OD_600_ measurement. The other half of the culture was used as the prey cells. Predator and prey suspensions were diluted to a final OD_600_ of 3 with 1X PBS. Subsequently, 4 µL of prey cell suspension were combined with 16 µL of predator cell suspension and subjected to gentle pulse-vortex to mix. 1 µL of such cell suspensions were applied in a 12 x 8 grid arrangement in a one-well plate containing 35 mL of LB agar. Up to 24 samples were spotted onto the agar in each one well plate. Samples were allowed to dry for 5 min at room temperature. Subsequently, the plate was incubated in a Biospa Automated Incubator (Biotek) at 30°C and the bioluminescence from prey cells was quantified over time using a Synergy Neo2 plate reader (Biotek). Under these assay conditions, maximal T6SS-driven killing of the Δ8 prey strain occurred at ∼125 min (Supplementary Figure 8). Data from this time-point are presented in the bar graphs in Figure 6A,C.

### Whole genome sequencing and variant calling

*V. cholerae* strains were diluted from freezer stocks into 3 mL of LB medium and cultured for 3-6 h until turbidity occurred (OD_600_ = 1-2). The cells were collected by centrifugation and DNA was purified from them using the DNeasy Blood & Tissue kit (Qiagen, Germany). Subsequently, the DNA was processed and sequenced. Variant calling to identify SNPs of interest was performed by the Microbial Genome Sequencing Centre (Pittsburgh, PA). *V. cholerae* N16961 was used as the reference genome.

### RNA isolation and quantitative RT-PCR

Strains were cultured for ∼18 h as described in the colony sectoring assay section. Subsequently, colonies were resuspended in 1X PBS, 4 mm glass beads were added, and the suspension subjected to vortex for 5 min to disperse aggregates. The resulting cell suspension was treated for 15 min at room temperature with RNAProtect reagent per the manufacturer’s instructions. Thereafter, RNA isolation, cDNA library preparation, and qPCR was performed as described previously (Mashruwala and Bassler, 2020).

### Image acquisition

#### Time-lapse acquisition

Colonies were plated as described in the colony sectoring assay section. Thereafter, images of growing colonies were acquired using a Cytation 7 imaging plate reader (Biotek) using the attached temperature-controlled incubator at 30°C and a 4× air objective. Live-cell distribution was monitored using intensity from a chromosomally-integrated fluorescent reporter that constitutively produced the mKO protein (ex: 500 nm). Dead-cell distribution was monitored using staining intensity from SytoX (ex: 556 nm). The focal plane was maintained using the laser autofocus method. For each time point and in each acquisition channel, a 3x3 *xy*-montage of the colony was obtained and stitched together using the linear blend algorithm to form a single image. In every case, a depth of 225 μm was sectioned. Maximum intensity *z*-projections were generated for each time point using the Biotek Gen5 software.

#### Bright-field and fluorescent stereo-microscope images

Colonies were plated as described above in the colony sectoring assay section. Following 2-3 days of growth, images were acquired using a Leica M125 stereo-microscope with a Leica MC170 HD camera.

### Image analyses and quantitation

Time-projections: Projections of time-lapse data were obtained using customized Fiji scripts that performed the following sequence of events: First, image background subtraction was performed using a rolling ball radius of 1,000 pixels. Second, to account for shifts during imaging, the sequence of images was registered using the MultiStackReg Fiji plugin and the Rigid Body algorithm. Next, the registered image sequences were collapsed using maximum intensity projections. Ratio images were obtained using the Fiji Image calculator tool to divide pixel intensities across the entire image of the dead-cell channel by that for the live-cell channel. The grayscale time-projections and ratio images were pseudo-colored using Red (live channel), Cyan (dead channel) or Mpl-inferno (ratio image) look-up tables. To aid in visualization, the time-projection images were cropped at the colony boundaries and pixel intensities outside the colony boundaries were set to zero.

Weka machine learning-dependent image segmentation: Time-projection images for the first 38 h of colony growth were segmented and analyzed using the Fiji Trainable Weka Segmentation tool. First, a classifier model was trained to discriminate between the sectored and non-sectored regions of the colonies. The training dataset consisted of 25 time-projection images from the dead-cell channel using images of both parent and mutant colonies from experiments performed on multiple days. Using a custom Fiji script, the resulting classifier model was applied to time-projection images to obtain probability images in which each pixel in an image was assigned a probability of belonging to a particular image class. Regions of interest (ROI) were extracted from these probability images by thresholding using the RenyiEntropy algorithm followed by application of a combination of the filter and the particle size cutoff tools which were customized for each segmentation class. The obtained ROIs were manually curated for mis-segmentation, and the curated ROIs were used in measurements of area or intensities from the time-projection or the ratio images.

Space-time kymographs: Fluorescence time-course images of colony growth were analyzed using a custom MATLAB script. First, the center of the colony was located with an iterative centroid-finding algorithm using the fluorescent channel that monitored live-cells, beginning at the first image acquisition at 8 h. To eliminate occasional sudden shifts due to mechanical noise, the sequence of images was registered in the *x*,*y* plane without rotation correction. Next, spatio-temporal fluorescence intensities in both the live- and dead-cell channels were extracted for kymograph analyses as follows: A region of interest consisting of a radial section, akin to a pie slice, was specified and manually verified to lack sectoring. Colony boundaries were determined using a fixed intensity threshold for the maximum fluorescence signals. Pixel intensities from both the live- and dead-cell channels were averaged in the circumferential direction within the radial section to obtain the averaged fluorescence intensity profile along the colony radius and over time. The obtained intensity values were used to construct kymograph profiles quantifying the space-time development of live and dead cells within the colony.

## Acknowledgements

We thank Wenjuan Du for the initial isolation of two of the variant strains as part of her undergraduate senior thesis research, and for creative and stimulating discussions. We thank Prof. Ned Wingreen for his generous feedback on our manuscript. We are grateful to members of the Bassler group for thoughtful discussion, especially Andrew Bridges for his generous help with instrument training. This work was supported by the Howard Hughes Medical Institute, National Science Foundation grant MCB-2043238, and NIH grant 5R37GM065859 to B.L.B. A.A.M. is a Howard Hughes Medical Institute Fellow of the Life Sciences Research Institute. The authors have no competing interests to declare.

## Supplementary Figures and Legends

**Supplementary Video 1:** T6SS activity drives two phases of spatio-temporal cell death in *V. cholerae* 2740-80, and Phase 1 precedes colony sectoring. Time-lapse image series for growth of *V. cholerae* 2740-80 constitutively producing mKO which marks live cells (left). Dead cells are marked with SytoX stain (middle). Intensity ratios (right) were obtained as in Figure 3. Intensities are color-mapped and the scale bars on the left of each panel represent color:intensity. Acquisition times are displayed on the bottom right of each panel.

**Supplementary Figure 1:**
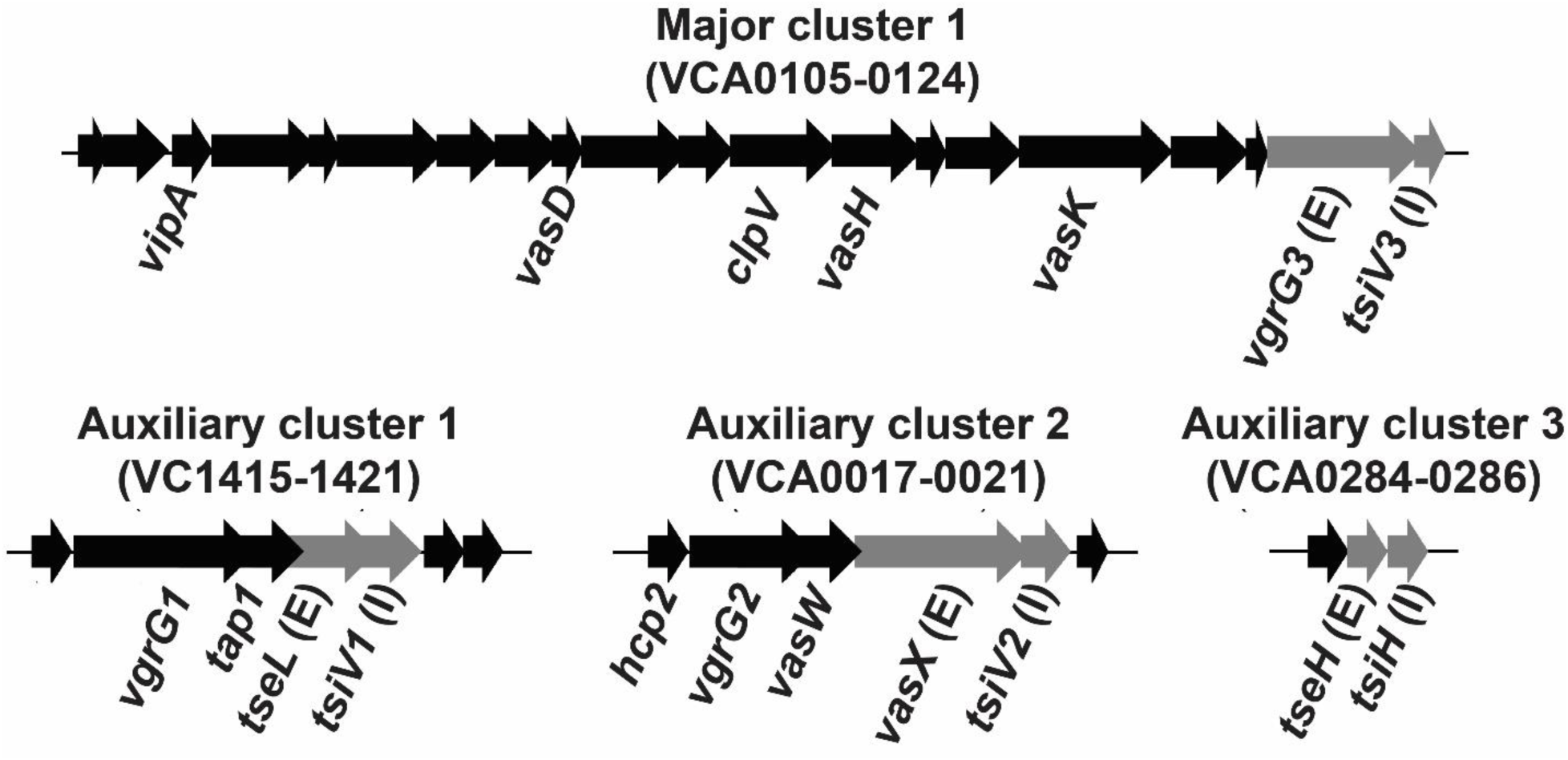
Arrangement of *V. cholerae t6ss* genes in four clusters. Select gene names are provided. Genes encoding effector and immunity proteins are depicted in gray and designated with, respectively, an E or I in parentheses (adapted from Metzger et. al, 2016).

**Supplementary Figure 2:**
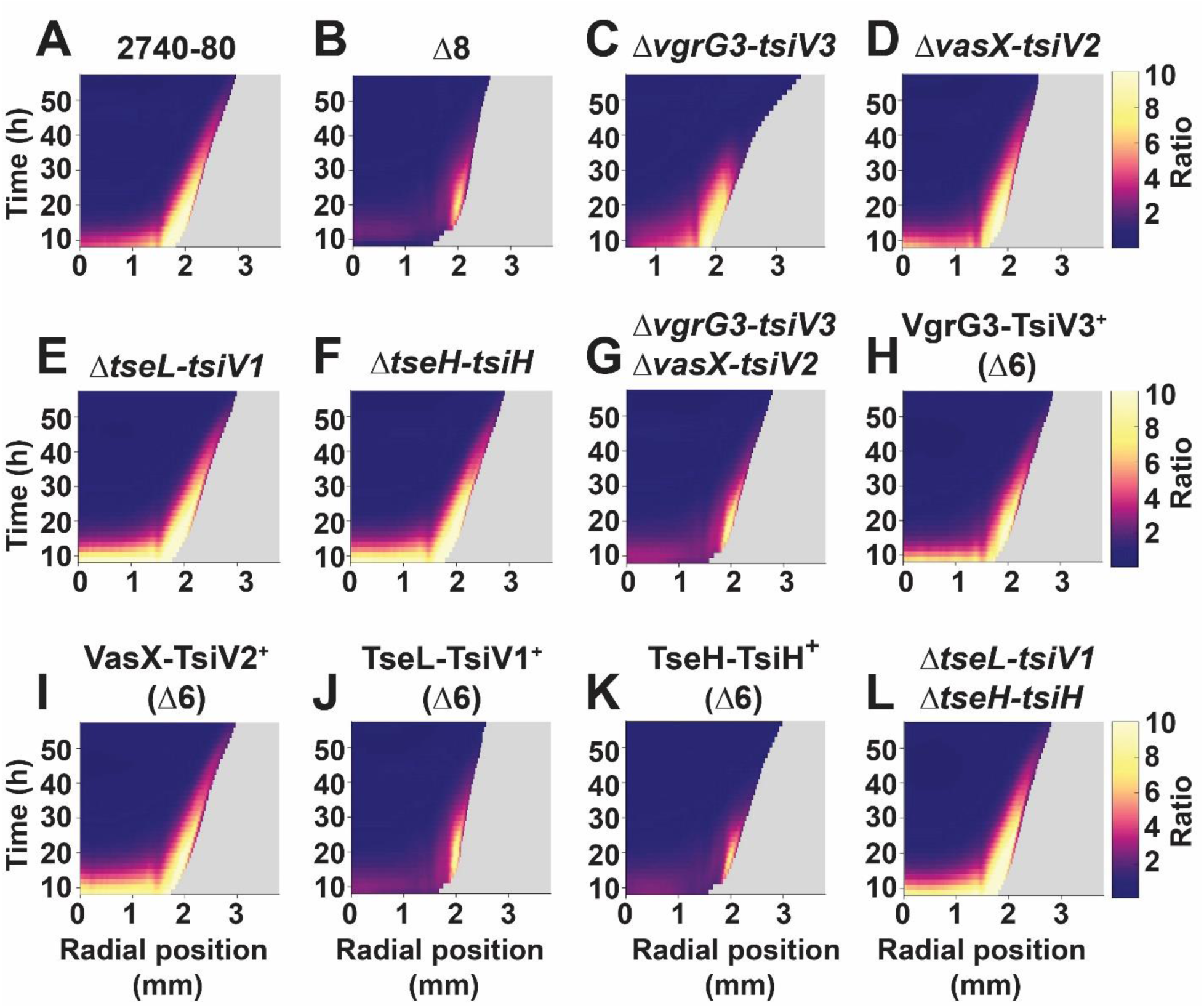
The VgrG3-TsiV3 and VasX-TsiV2 T6SS effector-immunity pairs drive Phase 1 cell death in *V. cholerae* 2740-80. Linear ratio kymographs for the indicated strains. These data accompany Figure 5. The linear scale emphasizes differences in Phase 1 cell death. Kymograph data treated as described for Figure 3. Kymographs from one colony are representative of results from 3-9 colonies for each strain. The Phase 1 cell death shown in panel C for the Δ*vgrg3-tsiv3* strain appears to end abruptly. This feature is due to a technical limitation. Specifically, the Δ*vgrg3-tsiv3* strain displays hyper-sectoring (see Supplementary Figure 4). Thus, by late Phase 1 (∼34-40 h), the outer regions of Δ*vgrg3-tsiv3* colonies are composed almost entirely of sectors. Because we exclude sectored regions from the kymograph analyses (see Methods), Phase 1 appears abbreviated in this strain, but that is not the case.

**Supplementary Figure 3:**
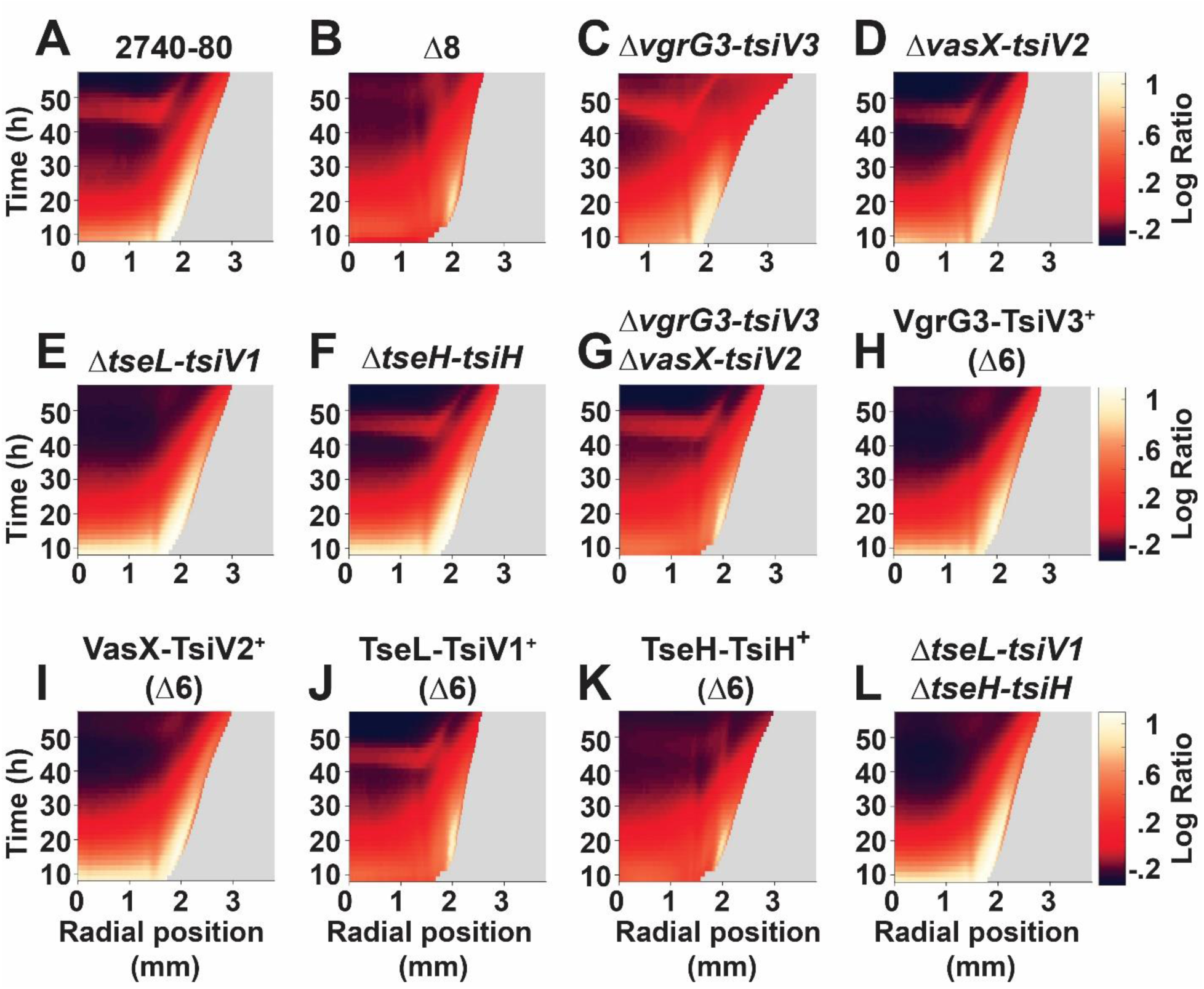
The TseH-TsiH effector-immunity pair is dispensable for T6SS-mediated cell death in *V. cholerae* 2740-80. Logarithmic ratio kymographs for the indicated strains. These data accompany Figure 5. Kymograph data treated as described for Figure 3. Kymographs from one colony are representative of results from 2-6 colonies for each strain. The abrupt truncation of Phase 1 cell death shown in panel C for the Δ*vgrg3-tsiv3* strain is a consequence of the imaging limitation in which sectors are omitted and is also described in the Supplementary Figure 2 legend. The order in which the strains are arranged is identical to that in Supplementary Figure 2.

**Supplementary Figure 4:**
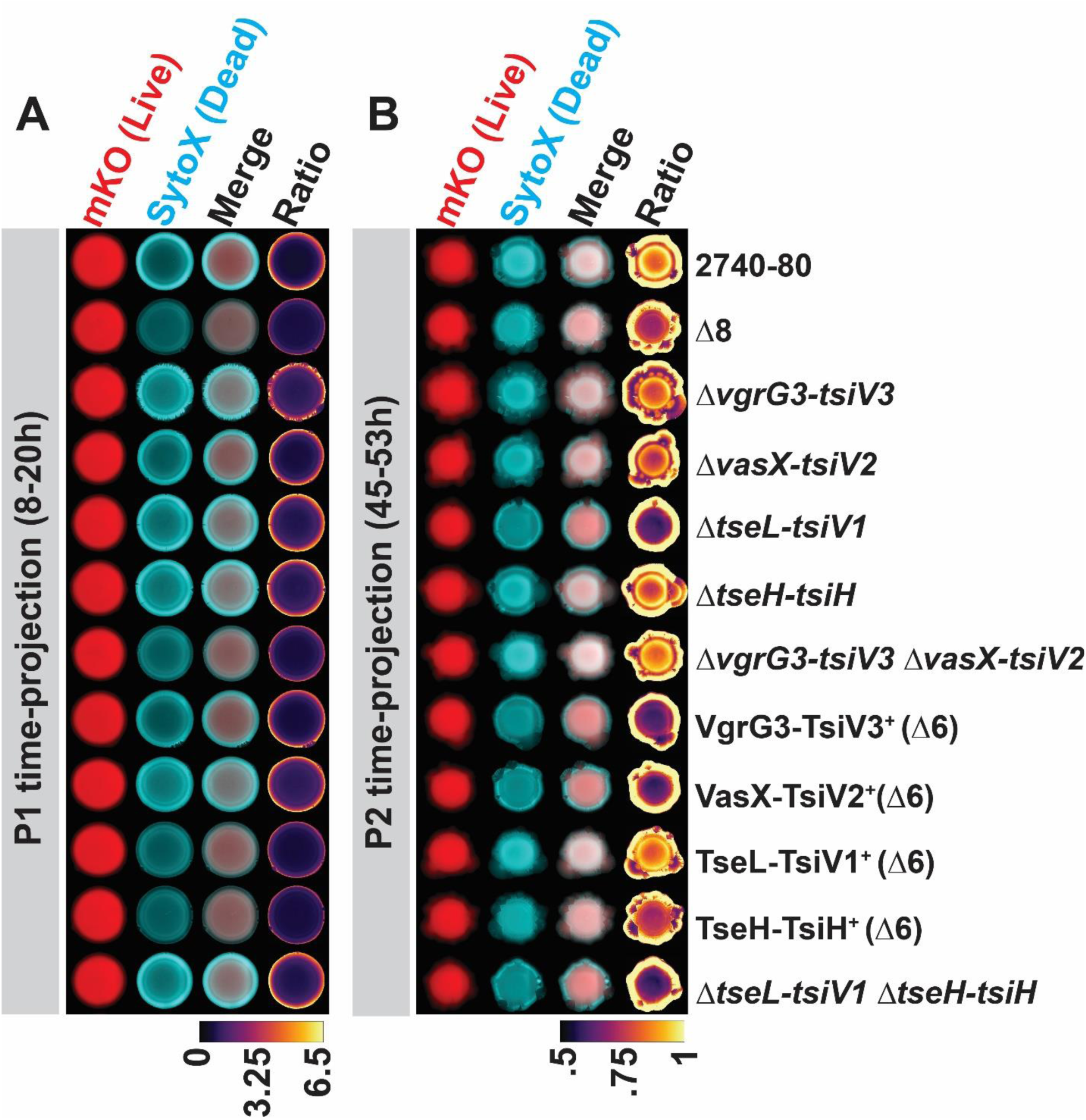
The VgrG3-TsiV3 and VasX-TsiV2 effector-immunity pairs drive Phase 1 cell death, while the TseL-TsiV1 effector-immunity pair drives Phase 2 cell death in *V. cholerae* 2740-80. Time-projections show cell death and sectoring for the indicated phases and strains. These data accompany Figure 5. The order in which the strains are arranged is identical to that in Supplementary Figures 2 and 3.

**Supplementary Figure 5:**
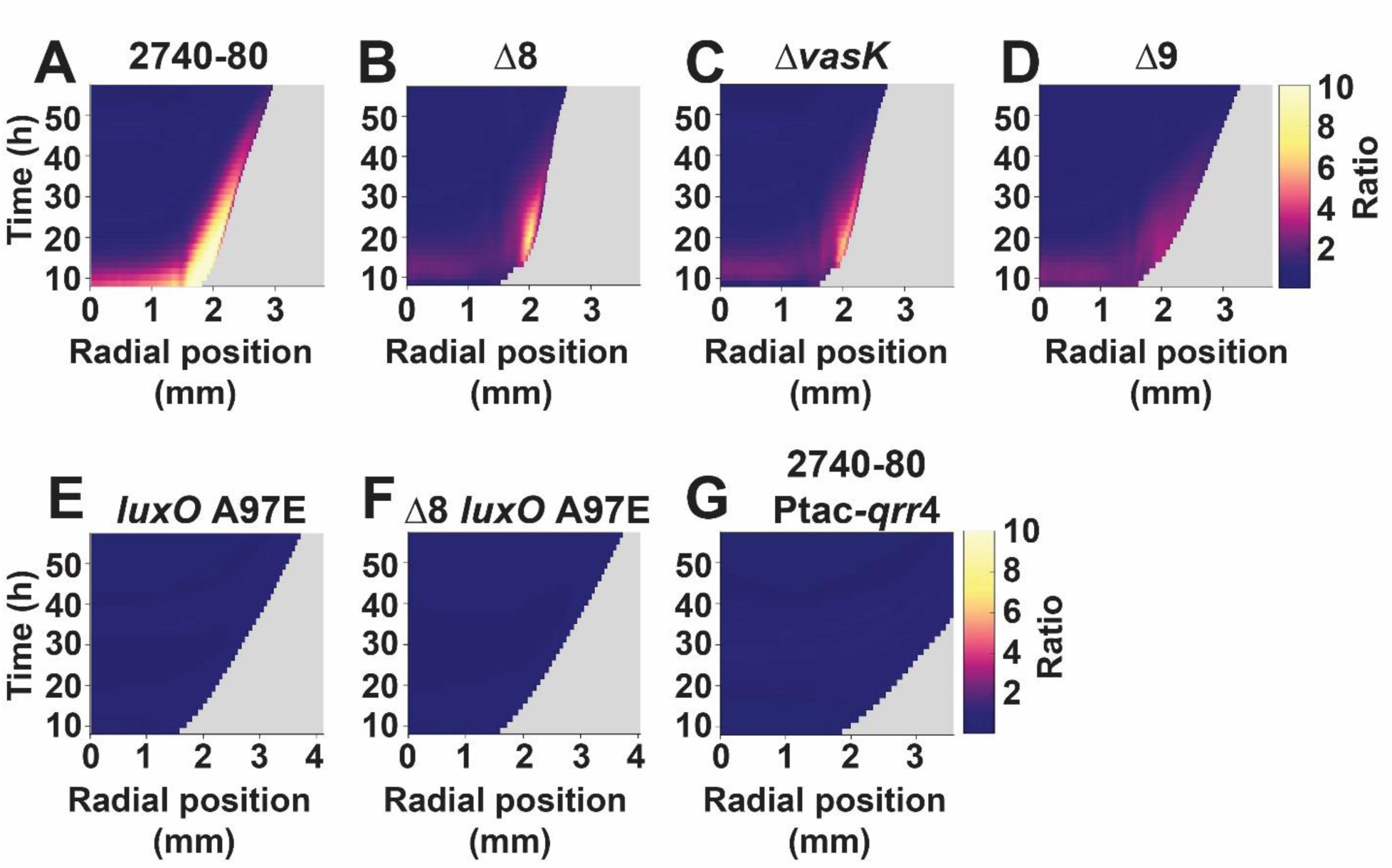
Strains lacking *vasK* or harboring *luxO* A97E are deficient in Phase 1 cell death. Linear ratio kymographs for the indicated strains. These data accompany Figure 5 and Figure 6. Kymograph data treated as described for Figure 3. Kymographs from one colony are representative of results from 3-9 colonies for each strain.

**Supplementary Figure 6:**
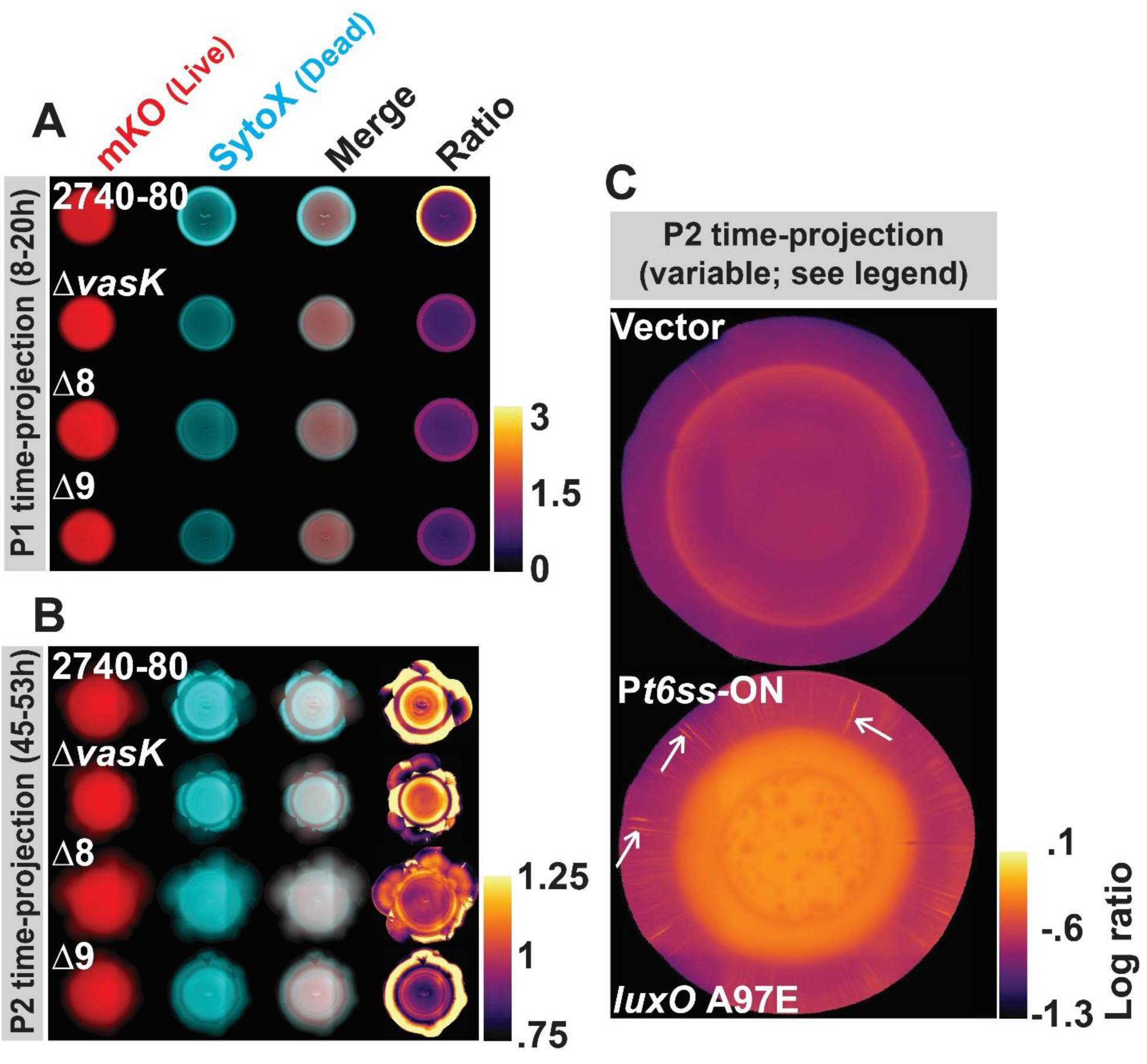
*V. cholerae* 2740-80 lacking all T6SS components displays less sectoring than *V. cholerae* 2740-80, the VasK-dependent T6SS-injection machinery is required for Phase 1 cell death but is dispensable for Phase 2 cell death, and production of T6SS components in a QS locked-LCD strain drives radial sectoring. (A-B) Time-projections showing cell death and sectoring for the indicated strains and phases. These data accompany those in Supplementary Figure 5A-D. (C) The *V. cholerae* 2740-80 *luxO* A97E strain carrying P*t6ss*-ON forms radial sectors. Shown are enlarged time-projection images of colonies of the indicated strains from Figure 7B. The white arrows point to radial sectors. Such sectors are not apparent in the strain carrying the empty vector.

**Supplementary Figure 7:**
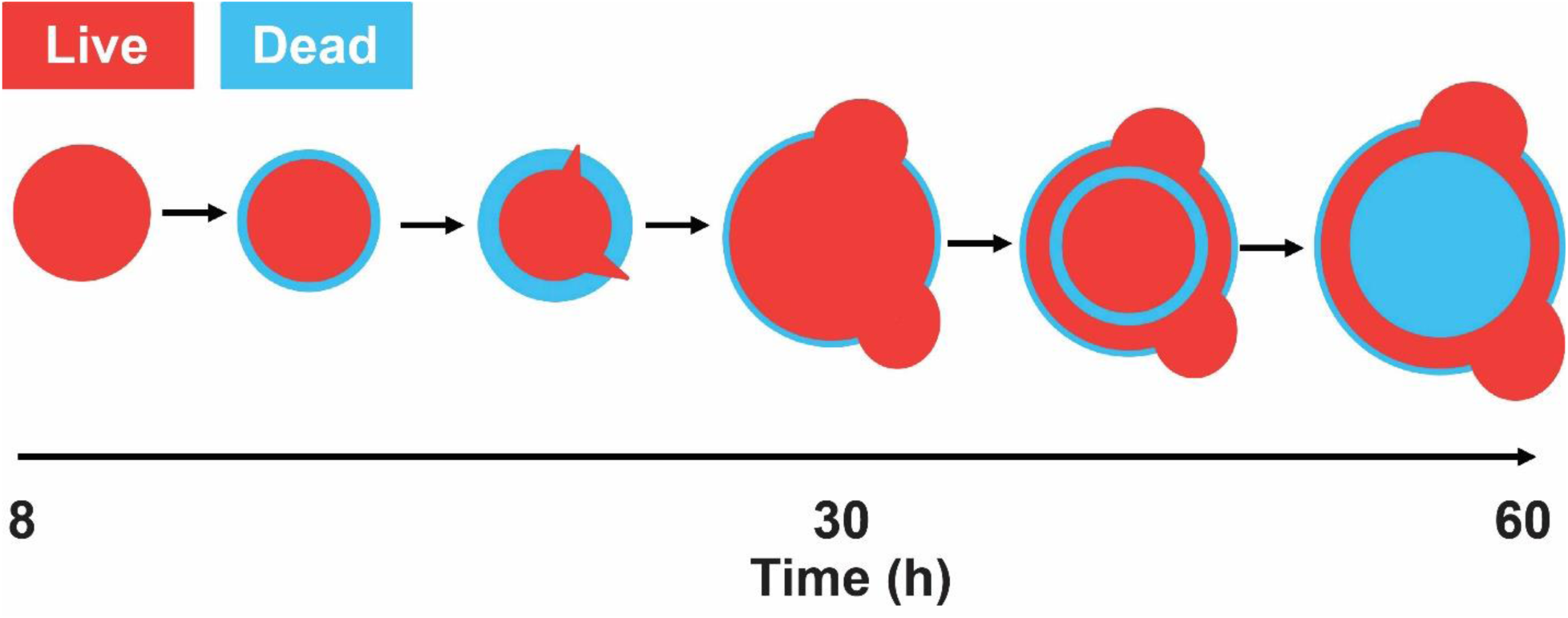
A model for cell death and sectoring in *V. cholerae* 2740-80 colonies. Colonies of *V. cholerae* strain 2740-80 undergo two-phase T6SS-mediated spatio-temporal cell death. Phase 1 initiates at ∼8 h post inoculation, occurs along the colony periphery, is driven by the VgrG3 and VasX toxins and is regulated by QS. At ∼18 h post inoculation, sectors emerge along the colony rim, i.e., from regions displaying high Phase 1 cell death. Cells in sectors undergo low cell death. Sectoring does not occur in a strain that is locked into the low cell density QS mode and that does not undergo Phase 1 cell death. Thus, Phase 1 cell death is key for sectoring to occur and, therefore, for enhanced genetic diversity to arise in the population. At ∼42 h, Phase 2 cell death initiates as a ring in the colony interior. Phase 2 cell death is driven by the TseL toxin and does not require the T6SS injection apparatus. Cell death propagates inward and outward from the initial ring.

**Supplementary Figure 8.**
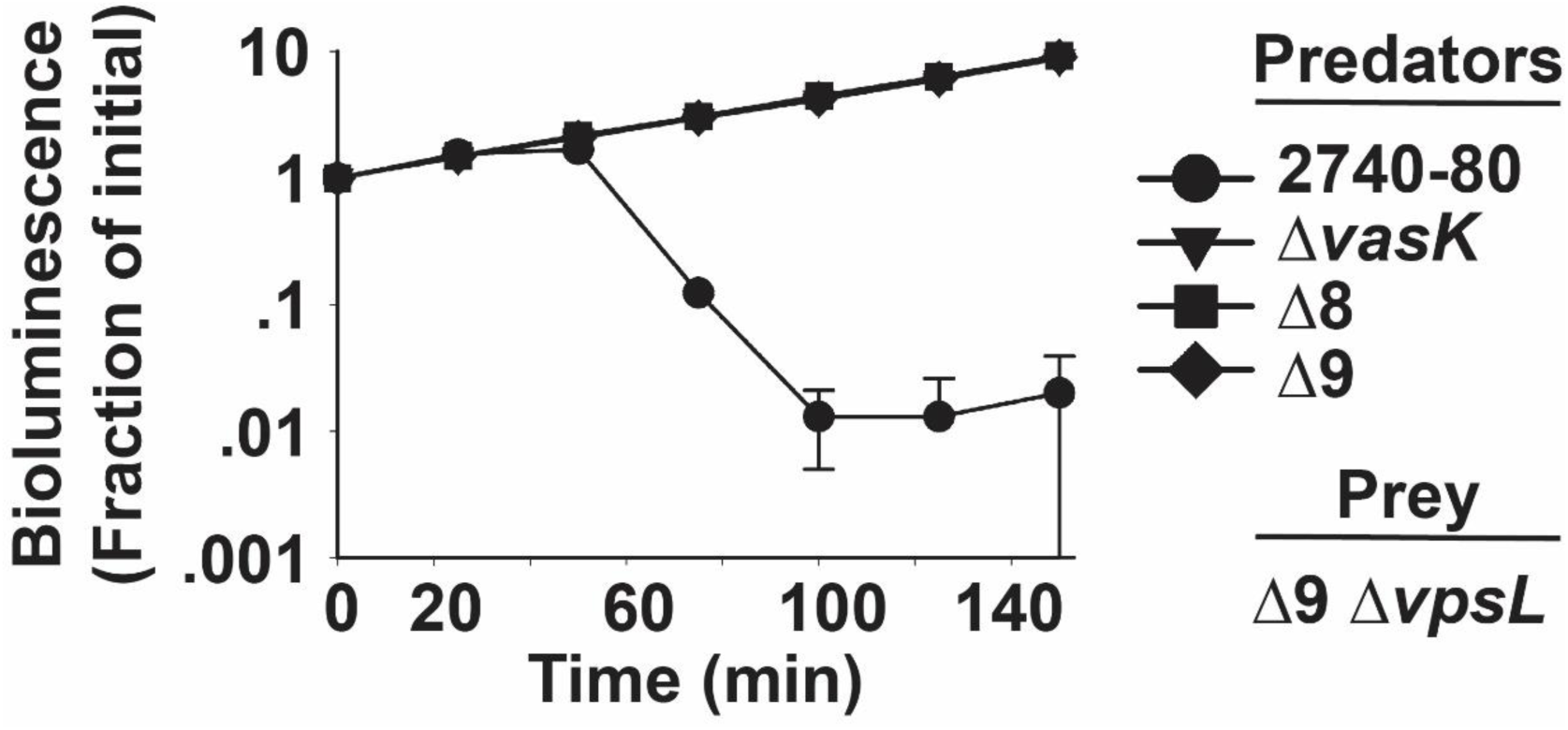
A functional T6SS apparatus is required for *V. cholerae* 2740-80 killing of bioluminescent T6SS-inactive prey cells. Inter-bacterial T6SS killing assay measuring time-dependent survival of *V. cholerae* 2740-80 Δ9 Δ*vpsL* prey cells following challenge with the indicated *V. cholerae* predators. The prey strain constitutively expresses luciferase and bioluminescence output was used to quantify live cells.

